# Biogeographic context mediates multifaceted diversity-productivity relationships in island and mainland forests

**DOI:** 10.1101/2023.07.17.549319

**Authors:** Maria Laura Tolmos, Nathaly R. Guerrero-Ramirez, Aitor Ameztegui, Martha Paola Barajas Barbosa, Dylan Craven, Holger Kreft

**Author notes:** Corresponding author: Maria Laura Tolmos; Biodiversity, Macroecology and Biogeography, University of Göttingen, Büsgenweg 1, 37077 Göttingen, Germany. Joint senior authors (Dylan Craven, Holger Kreft).

## Abstract

**Aim:** Growing evidence suggests that impacts of biodiversity loss on ecosystem functioning and nature’s contributions to people are usually negative, yet the magnitude and direction of these impacts can be variable across naturally-assembled ecosystems. A potential driver of variation in diversity-productivity relationships is the biogeographical context, which may alter these relationships *via* processes acting on the size and composition of the species pool like dispersal limitation, environmental filtering, speciation, and invasibility. However, the extent to which the relationships between biodiversity facets and forest productivity are shaped by the biogeographic context remains uncertain. Here, we examine the effects of taxonomic, phylogenetic, and functional tree diversity on aboveground productivity in climatically similar forests on islands and mainland.

**Location:** Continental and insular Spain.

**Time period:** 1997-2018.

**Major taxa studied:** Trees.

**Methods:** Using plot data from a national forest inventory, we assessed the influence of taxonomic, phylogenetic, and functional diversity on aboveground productivity using linear models and structural equation models, while accounting for environmental conditions, non-native species, and the number of trees.

**Results:** We find that drier environmental conditions lead to a decrease in productivity and in the number of trees in both island and mainland forests. In island forests, non-native species increased productivity directly and *via* their effects on phylogenetic diversity.

**Main conclusions:** Our results suggest that multifaceted diversity, by capturing the diversity of evolutionary history, contributes to elucidating diversity-productivity relationships in island forests that could not be detected otherwise by taxonomic diversity alone. By filling empty niches in island forests, we find that non-native species are fundamentally altering ecosystem functioning on islands.

## Introduction

There is compelling evidence that biodiversity positively influences ecosystem functioning across numerous experimental and real-world ecosystems (Cardinale et al., 2011; Flombaum & Sala, 2008; Gonzalez et al., 2020; Grace et al., 2016; Guerrero-Ramírez et al., 2017; Tilman et al., 2014). Specifically, positive effects of species coexistence through niche or resource partitioning and facilitation (Barry et al., 2019) promote higher ecosystem functioning through higher plant diversity and support the ability of more diverse plant communities to produce more biomass through complementarity (Cardinale et al., 2007; Hooper et al., 2005; Loreau & Hector, 2001). Alternatively, positive effects of plant diversity on biomass may be explained by more diverse plant communities having one or few species that are highly productive (Loreau & Hector, 2001). However, studies on the relationships between biodiversity and productivity in naturally-assembled forests have shown contrasting results across environmental gradients (Paquette et al., 2018; Ratcliffe et al., 2017; but see Liang et al., 2016). This suggests that elucidating biodiversity-ecosystem functioning (BEF) relationships in real-world ecosystems requires considering environmental conditions and - possibly - the biogeographic context in which these relationships occur.

Biotic and abiotic conditions have been found to strongly influence BEF relationships in forests (e. g. Craven et al., 2020; Fei et al., 2018; Jing et al., 2022; Mina et al., 2018; Ratcliffe et al., 2017), with forest types, geographic regions, and climatic conditions mediating the impacts of biodiversity on ecosystem functioning (Figure 1; Paquette & Messier, 2011; Pretzsch et al., 2013; Forrester, 2014; Grossiord et al., 2014; Jucker et al., 2016; Liang et al., 2016; Ratcliffe et al., 2016). For instance, water availability influences the strength of the BEF relationship in forests, with water-limited regions showing stronger positive BEF relationships than regions with higher water availability (Jing et al., 2022; Ratcliffe et al., 2017). Across latitudes, positive tree diversity-biomass relationships have been found consistently in temperate forests, while in tropical forests positive, negative, and neutral relationships have been observed (van der Plas, 2019). In addition, biogeographical context, i.e., the contrasting geographical locations, their geological histories, and their impact on shaping the processes generating biodiversity such as speciation and dispersal (Vellend, 2017), could be an important driver of the direction and magnitude of BEF relationships as they shape biodiversity patterns across spatiotemporal scales (e.g., Keil & Chase, 2019; Cai et al., 2023). Yet, the potential influence of biogeographic context on BEF relationships has rarely been assessed.

**Figure 1.**
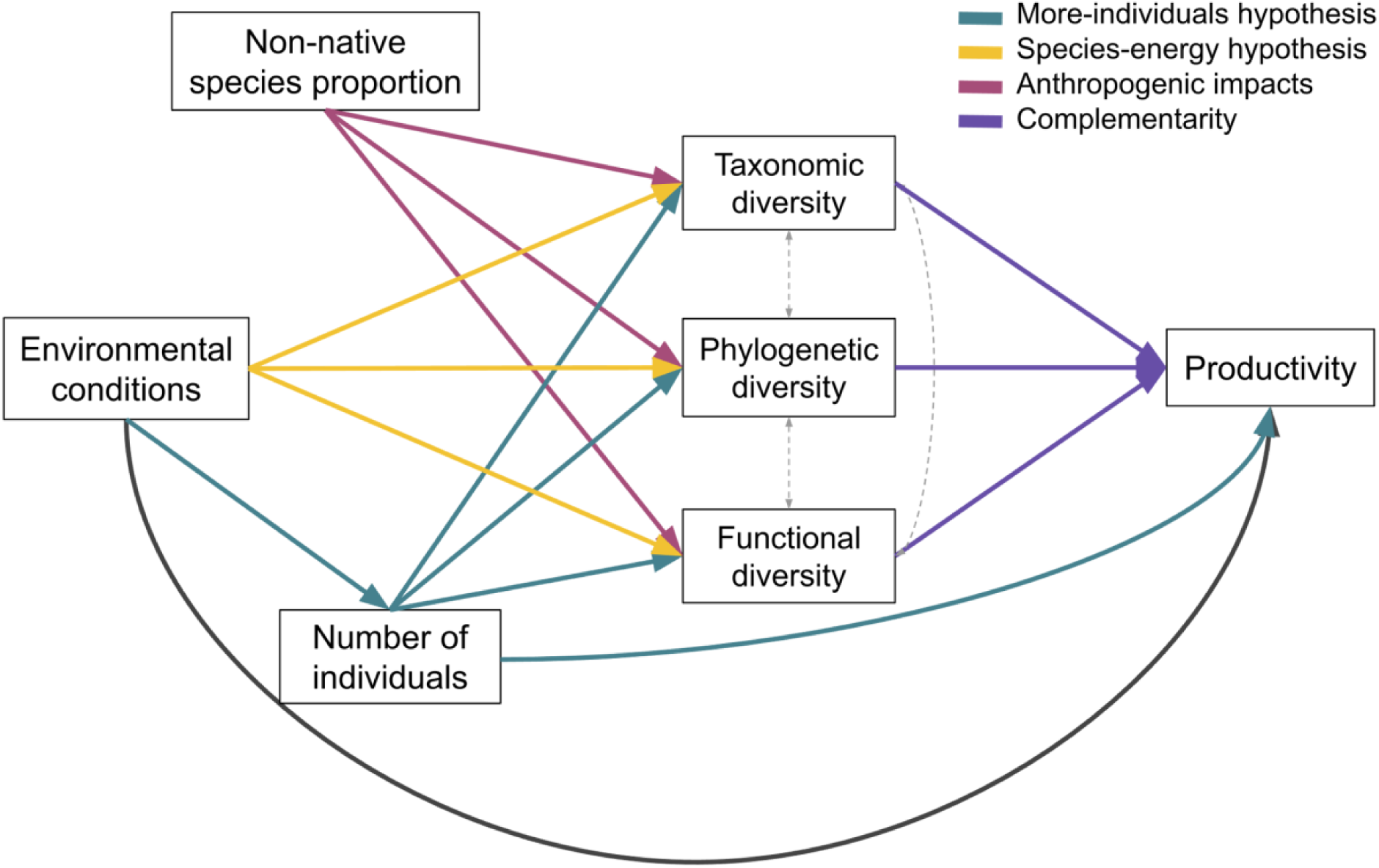
Conceptual model illustrating the expected effects of non-native species, environmental conditions, number of individuals, and multifaceted biodiversity on ecosystem functioning on mainland and island forests. Colored paths represent alternative possible ecological mechanisms and anthropogenic impacts influencing certain relationships. Black path is a relationship that could not be linked to a specific theoretical expectation. While we expected these pathways to influence overall relationships in both mainland and island forests, we hypothesized a stronger BEF relationship in islands due to their evolutionary history, functional adaptations, and vulnerability to invasion. Grey dashed arrows are expected correlations between variables.

Theories that attempt to disentangle biodiversity spatial patterns may contribute directly or indirectly to understanding biodiversity effects on ecosystem functioning. For instance, the more-individuals hypothesis proposes that the total number of individuals in a community limits the number of species with viable populations, with a higher number of individuals resulting in higher species richness via passive sampling and therefore potentially higher productivity (Figure 1; Gaston, 2000; Srivastava & Lawton, 1998). Conversely, the species-energy hypothesis proposes that resource availability, e.g., temperature, water availability, limits population sizes in a given area, regulating the number of individuals, species richness and, consequently, the capacity of a given community to produce biomass via niche-based processes (Figure 1; Brown, 2014; Wright, 1983).

Early BEF studies focused on how species richness drives ecosystem functioning (van der Plas, 2019) and largely ignored the influence of species abundances, evolutionary history, and functional differences among species. Additionally, most studies in naturally-assembled ecosystems investigate taxonomic diversity, despite the potential influence of other biodiversity facets on ecosystem functioning (Hagan et al., 2023; van der Plas, 2019). By embracing the multifaceted nature of biodiversity, we can examine ecological and evolutionary processes shaping species assemblages and diversity patterns beyond species counts, allowing us to better understand the impact of biodiversity change on ecosystem functioning (Cadotte et al., 2011; Díaz et al., 2007; Emerson & Gillespie, 2008). Specifically, functional diversity, *via* niche complementarity, not diversity *per se* influences ecosystem functioning (Díaz & Cabido, 2001; Flynn et al., 2009; Loreau, 1998; Tilman et al., 1997). Further, phylogenetic diversity may explain variation in ecosystem functioning (Cadotte et al., 2008; Venail et al., 2015) by representing the diversity of phylogenetically conserved functional traits and integrating a greater number of traits than the soft ones usually used to estimate functional diversity (Nock et al., 2016). However, biodiversity-productivity relationships may be driven by functional traits that are not phylogenetically conserved, and would therefore be harder to capture by phylogenetic diversity indices alone (Craven et al., 2018), highlighting the importance of including additional facets of diversity (Figure 1).

Their limited area, varying levels of isolation, and contrasting geological histories make oceanic islands perfect study systems to understand how different ecological and evolutionary processes shape diversity patterns and species assemblages, and how biogeographical context influences them (Hagan et al., 2021; Warren et al., 2015; Weigelt et al., 2015; Whittaker & Fernández-Palacios, 2007). Processes such as rare dispersal events, environmental filtering, and in-situ speciation have generated unique, highly endemic plant assemblages on islands worldwide (Weigelt et al., 2015). The biased representation of higher taxa compared to the source pool on islands due to dispersal, environmental, and biotic filters, i.e., disharmony, (Carlquist, 1974; König et al., 2021; Kraft et al., 2015) may result in closely-related species occupying different ecological niches, potentially increasing ecosystem functioning. Yet, unfilled niche space may persist on oceanic islands due to limited colonization and low species diversity. Further, the high proportion of endemic species makes island ecosystems more susceptible to the naturalization of non-native species (Moser et al., 2018; Sax et al., 2002), particularly on islands where phylogenetic relatedness among native species is higher (Bach et al., 2022). Moreover, there is a trend of disproportionate losses of island endemic species, as around 60% of all recorded extinctions took place on islands (Whittaker & Fernández-Palacios, 2007).

Their low species richness and high invasibility makes island ecosystems sensitive to changes in the abundance of keystone species, with potential repercussions on nutrient cycling, primary productivity, and carbon storage (Worm & Duffy, 2003). In addition, anthropogenic impacts on island biodiversity and ecosystem functioning are a growing concern. Non-native species - particularly invasive species - and land-use change can modify ecosystem structure and function by negatively impacting native island ecosystems through changes in nutrient cycling, carbon storage, altering species composition, and increasing fire risk (Figure 1; Mascaro et al., 2012; Rothstein et al., 2004; Vitousek et al., 1996; Vitousek et al., 1997). However, non-native species may also increase certain ecosystem functions in highly degraded islands, enhancing soil structure and fertility and restoring forest cover (Lugo, 2004). Similarly, there is evidence that novel forests, i.e forests resulting from a mixture of native and non-native species, can be more diverse and provide higher levels of ecosystem functioning than uninvaded native forests (Mascaro et al., 2012).

Here, we examine the relationship between taxonomic, phylogenetic, and functional diversity and aboveground productivity, an important ecosystem function, in forests on the Canary Islands and climatically similar areas in mainland Spain, and the extent to which multifaceted BEF relationships are mediated by biogeographical context. To this end, we used forest inventory data from Spain to first examine a simple version of the biodiversity-productivity relationship, i.e., not accounting for environmental conditions, for island and mainland forests. Later, we included the direct and indirect influence of environmental conditions and ecological processes that may affect this relationship (Figure 1). Specifically, we expected (1) positive multifaceted biodiversity-productivity relationships due to complementarity acting on both island and mainland forests, with phylogenetic diversity having a stronger influence on productivity than taxonomic or functional diversity as it can estimate the functional trait space of a community and also reflect species interactions (Srivastava et al., 2012), and phylogenetic diversity has been found to promote ecosystem functions and stability (Cadotte et al., 2012; van der Plas, 2019; Venail et al., 2015). We hypothesized that the magnitude of multifaceted biodiversity-productivity relationships differ between biogeographic contexts, with stronger relationships on islands because a large proportion of island biota evolved in these ecosystems and therefore developed specific traits to more efficiently use limited resources or to persist in harsh environments (Barajas Barbosa et al., 2023; Emerson & Gillespie, 2008); (2) environmental conditions, i.e., climate and soil properties, influence productivity on island and mainland forests, as bioclimatic variables and soil nutrients have been shown to strongly influence plant diversity (Kreft & Jetz, 2007; Lambers et al., 2011) and productivity (following the species-energy hypothesis); (3) the number of individuals positively influences productivity on island and mainland forests, as a higher number of individuals is expected to host a more biodiverse community (following the more-individuals hypothesis), which subsequently is expected to yield higher productivity (Gaston, 2000; Srivastava & Lawton, 1998); and (4) non-native species potentially influence productivity in both mainland and island forests, with the effect being stronger and positive on islands as non-native species likely perform different functions than native species (Rothstein et al., 2004; Vitousek et al., 1996). However, as the Canary Islands forests may not have been impacted by non-native species as extensively as other oceanic islands (Fernández-Palacios et al., 2023), the influence of non-native species on ecosystem functioning may be similar to that of mainland forests.

## Methods

### Study area

Our main goal was to study the relationship between multiple diversity facets and forest productivity in two areas with similar climates but contrasting biogeographical contexts. We selected the Canary Islands archipelago (hereafter ‘the Canaries’) as our focal oceanic archipelago because its flora has been extensively studied (Acebes-Ginovés et al., 2010; Barajas Barbosa et al., 2023) and it has broad environmental gradients in elevation and aridity (Ashmole & Ashmole, 2016; Del Arco Aguilar & Rodríguez Delgado, 2018). Located in the Atlantic Ocean off the northwestern tip of Africa, the Canaries are an archipelago of volcanic origin formed progressively starting around 60 Ma (Troll & Carracedo, 2016). Island age increases from west to east, with the youngest island being El Hierro (1 Ma) and the oldest Fuerteventura and Lanzarote (ca. 25 Ma), while elevation varies as a result of their geological development (Troll & Carracedo, 2016). Forested areas in the western Canaries are characterized by the Laurel Forest (or laurisilva) and the Canary Pine Forest. The laurisilva is dominated by the Lauraceae family and occurs on the windward (northern) side of the slope, linked to the presence of trade-wind clouds that help maintain the water balance by reducing evapotranspiration (Del Arco Aguilar & Rodríguez Delgado, 2018). The Canary Pine Forest is dominated by *Pinus canariensis,* which grows in a variety of habitats, most commonly between 500 to 1500 m a.s.l. on rocky outcrops, and on volcanic soils in areas with good drainage. Further, *P. canariensis* is a drought tolerant and fire resistant species (Santos et al., 2011).

We selected climatically similar regions on the Spanish mainland based on the Köppen-Geiger climate classification maps (Beck et al., 2018; Köppen, 1884). We restricted our analysis only to forests in areas that fall within the Köppen-Geiger category “temperate, dry summer, warm summer” (Csb), as this was the main category present in forested areas in the Canaries. Due to the greater climatic similarity between continental Spain and the western Canaries, we focused our study on the following islands: Tenerife, La Gomera, El Hierro, La Palma, and Gran Canaria. The selected mainland regions are located in the northwestern (Galicia) and northeastern (Catalonia) tips of continental Spain. Galicia is characterized by an oceanic climate, strongly influenced by the humid winds from the Atlantic with mild temperatures and abundant rainfall throughout the year. Galician forests are dominated mainly by temperate deciduous species such as *Quercus robur*, *Quercus pyrenaica*, and *Castanea sativa*, and *Pinus pinaster* forests towards the interior. Catalonia has a more Mediterranean to sub-Mediterranean character. Vegetation is dominated by pines (e.g., *Pinus halepensis*, *Pinus nigra* and *Pinus sylvestris*) and sclerophyllous hardwoods, particularly *Quercus ilex*.

### Data preparation

#### Forest composition and distribution

To assess aboveground productivity in naturally-assembled forests, we used the third (1997-2007) and fourth (2008-2018) Spanish National Forest Inventories (hereafter ‘forest inventories’). Forest inventories plots consist of nested concentric circular plots of 5, 10, 15, and 25 m radii based on DBH classes (for more details see: Dirección General para la Biodiversidad, 2007). We limited our analysis to plots occurring in forests without silvicultural treatments that were sampled in both forest inventories at the same location. Our study comprises 2,072 plots of 1963.5 m^2^ in continental (1434 plots) and insular Spain (637 plots; Figure 2b and c). In each plot, all trees with a diameter at breast height (DBH) ≥ 7.5 cm were recorded and identified, and their total height and DBH were measured (Alberdi et al., 2017).

**Figure 2.**
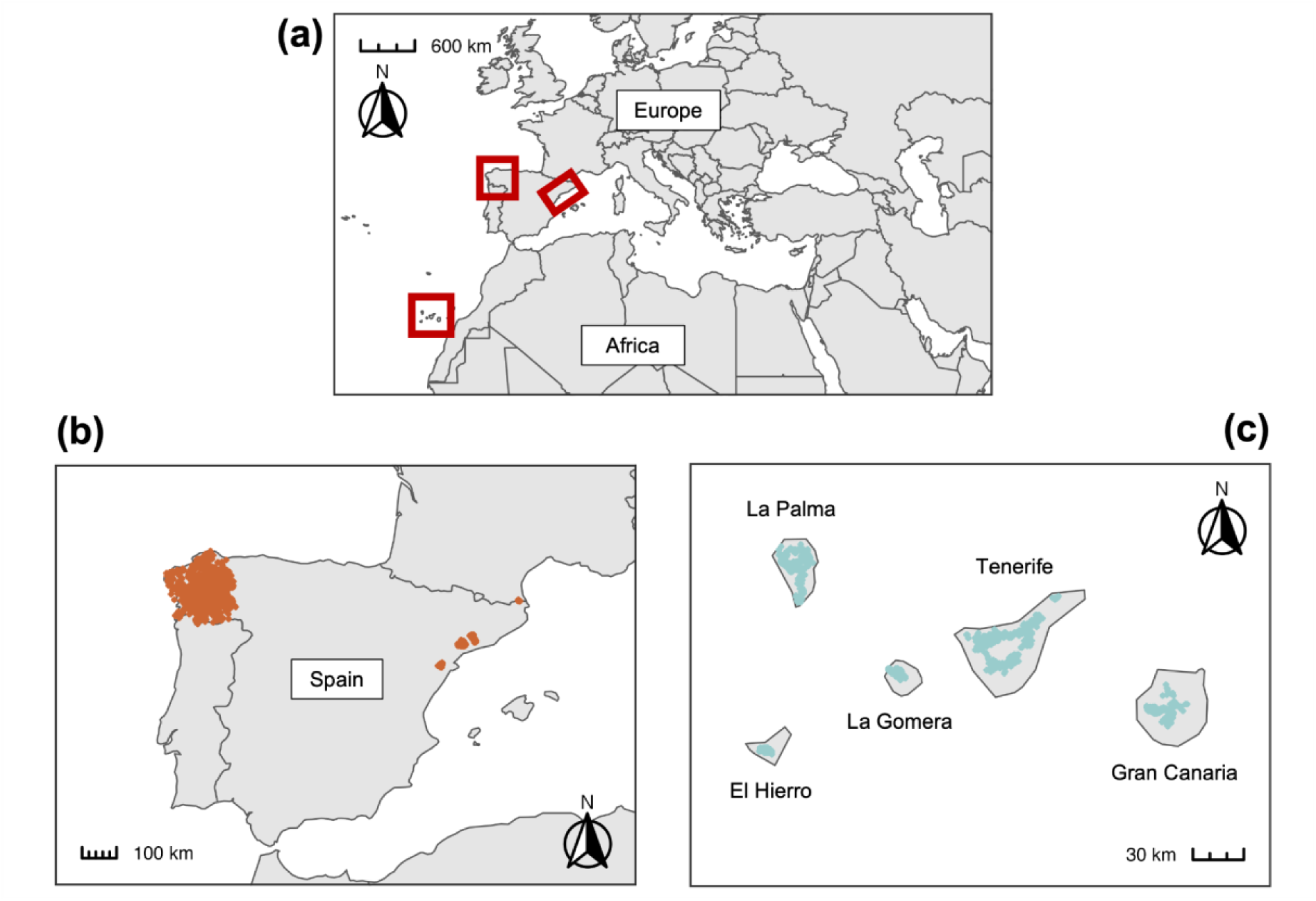
(a) Map of the study area in the Canary Islands and continental Spain indicated in red boxes, and the distribution of the study plots in mainland (b) and island (c) forests in regions classified as “temperate, dry summer, warm summer” (Csb) under the Köppen-Geiger climate classification.

#### Taxonomic standardization

Species names were standardized based on the Taxonomic Name Resolution Service version 5.0 (Boyle et al., 2013), Tropicos (Missouri Botanical Garden, 2021), and the World Checklist of Vascular Plants version 2.0 (WCVP, 2021) using the *TNRS* function of the R package ‘TNRS’ (Boyle et al., 2021). After name standardization, our dataset contained a total of 26 and 72 accepted tree species for the Canaries and mainland Spain, respectively (Table S1).

#### Phylogeny

We constructed a phylogenetic tree for all tree species using the seed plant phylogeny of Smith and Brown (2018) as a backbone, and subsequently added species to the backbone using dating information from congeners in the tree with the *congeneric.merge* function of the R package ‘pez’ (Pearse et al., 2015). All mainland and island species were successfully placed in the phylogeny.

#### Functional traits

To capture plant acquisitive strategies (Díaz et al., 2016; Reich, 2014; Wright et al., 2004), we obtained data for four functional traits: (1) maximum height (H; m) captures light interception ability (Moles et al., 2009; Westoby et al., 2002); (2) specific leaf area (SLA; mm^2^/mg) relates to carbon gain and leaf lifespan (Wright et al., 2004); (3) leaf area (LA; mm^2^) is important for light interception and influences leaf energy and water balance (Farquhar et al., 2002; Givnish, 1987); and (4) leaf nitrogen content per leaf dry mass (N_mass_; mg/g) influences leaf photosynthetic capacity, where higher values are correlated with higher photosynthetic potential (Díaz et al., 2016). These traits were measured locally for all native species in Tenerife using standard protocol following Pérez-Harguindeguy et al. (2013) and were obtained from Barajas Barbosa et al. (2023). For the remaining 15 island and 72 mainland species, we obtained functional trait data from TRY (Kattge et al., 2020). We restricted our analysis to 1813 plots (628 in insular and 1185 in mainland Spain) for which we had a minimum of 80% trait data coverage per plot (Pakeman & Quested, 2007). Our final trait dataset contained a total of 25 and 70 species for the Canaries and mainland Spain, respectively, but not all species had data for all four traits. For the 25 species in the Canaries, 84% of the species had trait data for H, SLA, and N_mass_, and 80% for LA. While for the 70 species in mainland Spain, 94.3% had available data for H, 92.9% for SLA, 84.3% for LA, and 91.4% for N_mass_. We performed phylogenetic trait imputation for the remaining missing data using the random forest algorithm with the *missForest* function of the ‘missForest’ package (Johnson et al., 2021; Stekhoven & Buehlmann, 2012; Figure S1). We then selected the number of phylogenetic eigenvectors (n=1-69 in mainland, n=1-24 in islands) that minimized the imputation error for each trait (Tables S2 & S3).

#### Environmental variables

We selected mean annual temperature (°C), mean annual precipitation (mm/a), potential evapotranspiration (PET; mm/a), aridity, wind speed (m/s), temperature seasonality (°C), and precipitation seasonality (mm) as bioclimatic variables as these regulate energy and water availability for plants and therefore have a strong influence on plant diversity and productivity (Kreft & Jetz, 2007). All bioclimatic variables, except for PET and aridity, were derived from the Climatologies at high resolution for the earth’s land surface areas (CHELSA V2.1; Karger et al., 2017). PET was derived from the Global Aridity Index dataset based on the FAO Penman-Monteith equation (Trabucco & Zomer, 2018), while aridity was calculated as mean annual precipitation/PET (lower values indicate drier conditions). Additionally, we derived soil nitrogen (cg/kg) and soil pH at a 15-30 cm depth from SoilGrids (Batjes et al., 2020) as soil nutrient availability has been shown to affect plant diversity (Lambers et al., 2011). It is important to mention that while study sites in mainland and insular Spain fall into the same Köppen-Geiger climate classification, moderate differences in the range of climatic conditions exist (Figures S2a-i and S4b).

#### Non-native tree species

We calculated the proportion of non-native tree species as the total number of non-native individuals per plot divided by the total number of individuals per plot (Figure S2j). Non-native tree species were determined based on species checklists from the GIFT database (BioScripts, 2014; Euro+Med, 2006; Rhind, 2020; Weigelt et al., 2020).

#### Number of individuals

We estimated the number of individuals as the average number of tree individuals for each plot in both forest inventories (Figure S2k).

### Ecosystem functioning

We used the annual volume increment as a proxy for productivity, one of the most widely studied ecosystem functions (Paquette & Messier, 2011; Craven et al., 2020). Volume values were calculated as part of each forest inventory using DBH and height measurements in combination with species-specific allometric equations fitted for Spanish forests (Dirección General para la Biodiversidad, 2007). Volume values in the forest inventories were calculated for each tree in a plot and extrapolated to an area of one hectare. To allow for comparisons with diversity metrics (see Simons et al., 2021), we interpolated volume values to the largest plot size. Finally, we estimated volume increment per plot as the difference between volume in the third and fourth forest inventories in m^3^.

### Multifaceted diversity

We estimated diversity metrics for each plot based on the framework by Chao et al. (2014), to make the three facets of diversity more comparable and to detect if diversity patterns are driven by rare or more common species in the assemblages. Species abundances of individuals that belonged to smaller DBH classes were linearly extrapolated to the largest plot size in order to make diversity metrics and annual volume increment comparable. However, as the species-area relationship is non-linear, this could bias diversity metrics. For *taxonomic diversity*, all species are taxonomically equally distinct, and the taxonomic entities represent species. We estimated taxonomic diversity as the effective number of species (Jost, 2006). Where the effective number of species is of order zero (q=0, ^0^D), it is the equivalent of species richness and does not take species abundances into account and is sensitive to sampling bias; in contrast, where the effective number of abundant species is of order two (q=2, ^2^D), it emphasizes abundant species and discounts rare ones (Chao et al., 2014). We estimated phylogenetic diversity as the effective number of phylogenetic entities (Chao et al., 2014), where a phylogenetic entity represents a unit length of a branch segment, and each segment is equally distinct phylogenetically. When the effective number of phylogenetic entities is of order zero (q=0, ^0^D), it is equivalent to Faith’s PD (Faith, 1992) and does not account for abundances, and order two (q=2,^2^D), as with taxonomic diversity, gives more weight to abundant phylogenetic entities (Chao et al., 2010). Finally, for *functional diversity*, the functional entity represents a unit of functional trait distance between two species, and all entities are equally distinct functionally. We estimated the effective number of functional entities (i.e., effective sum of functional distances between two species) (Chao et al., 2014). Where the effective number of functional entities is of order zero (q=0,^0^D), it is the equivalent of FAD (Functional Attribute Diversity) and, as with the previous two diversity facets, it does not take abundances into account, while the effective number of functional entities of order two (q=2,^2^D) favors abundant species (Chiu & Chao, 2014). All diversity indices were calculated using the R package ‘hillR’ (Li, 2018).

### Statistical analysis

To examine the influence of biodiversity on productivity, we performed our analysis in two steps. First, we tested the influence of biodiversity on productivity using linear models for each diversity facet and Hill number, including an interaction term between biogeographic context (mainland or island) and diversity facet (Figure 4, Table S4). We evaluated multicollinearity by calculating the variance inflation factor (VIF) for each model (Fox & Monette, 1992). We then tested the models for spatial autocorrelation with Moran’s I statistic, and found no significant spatial autocorrelation for any of the models (Moran’s I < 0.3). Model assumptions were checked visually for all models. F tests were calculated for all models with the *Anova* function in the R package ‘car’ (Fox & Weisberg, 2019). We visualized the relationship between explanatory variables and productivity using the R package ‘ggeffects’ (Lüdecke, 2018). Second, to assess the direct and indirect influence of environmental conditions, number of individuals, non-native species, and diversity facets on productivity, we used piecewise structural equation models (SEMs; Lefcheck, 2016). SEMs allowed us to integrate multiple response and predictor variables in an integrative model, in order to acquire a deeper understanding of the relationships between productivity and the variables that influence it, as well as the relationships between environmental conditions, non-native species, number of individuals, and diversity facets (Figure 1). Taxonomic and phylogenetic diversity indices, wind velocity, aridity, soil nitrogen, soil pH, and number of individuals were log_10_-transformed due to skewed distributions. All variables were further standardized using a z-transformation to make model coefficients comparable. Due to multiple environmental variables being highly correlated (Figure S3), we used the first axis (PC1) of a principal component analysis (PCA; Figure S4, Table S5). PC1 explained 57.9% of total variation and was mostly associated with water availability-related variables and soil pH. We tested SEMs paths for mainland and island plots separately for Hill numbers 0 and 2 based on directed separation tests. For mainland forests, we included direct paths between i) productivity, diversity facets, environmental variables, number of individuals, and non-native tree species; ii) each diversity facet, environmental variables, number of individuals, and non-native tree species; and iii) number of individuals and environmental variables. For island forests we excluded direct paths between taxonomic and functional diversity and productivity, and added a direct path between non-native species proportion and productivity. Fisher’s C statistics were calculated for all models, where p-values > 0.05 represented a good fit of the data to the hypothetical causal model (Lefcheck, 2016). We used the R packages ‘piecewiseSEM’ to calculate all SEMs (Figure 5; Lefcheck, 2016) and ‘semEff’ to calculate direct, indirect, total, and mediator effects on productivity (Tables S10-S13; Murphy, 2022) .

**Figure 4.**
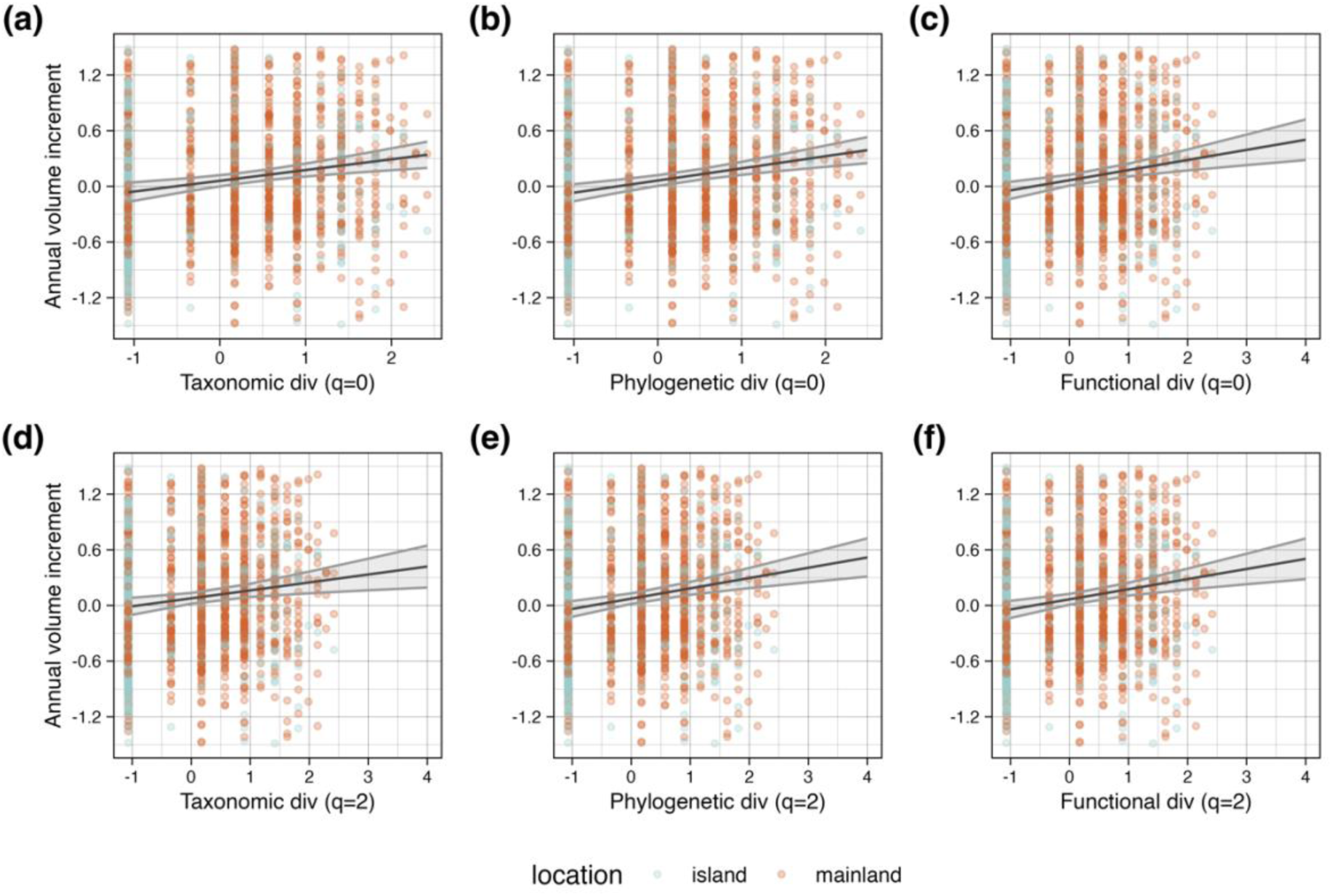
Linear models examining the separate influence of taxonomic (a,d), phylogenetic (b,e), and functional (c,f) diversity on productivity without considering environmental conditions in mainland and island forests in Spain in regions classified as “temperate, dry summer, warm summer” (Csb) under the Köppen-Geiger climate classification. Annual volume increment is used as a proxy for productivity. Separate models were fitted for Hill numbers 0 (i.e., species richness; a-c) and 2 (d-f). All variables are scaled. All predicted values are significant (p-value < 0.01). Dots represent data points for islands (turquoise) and mainland (orange).

**Figure 5.**
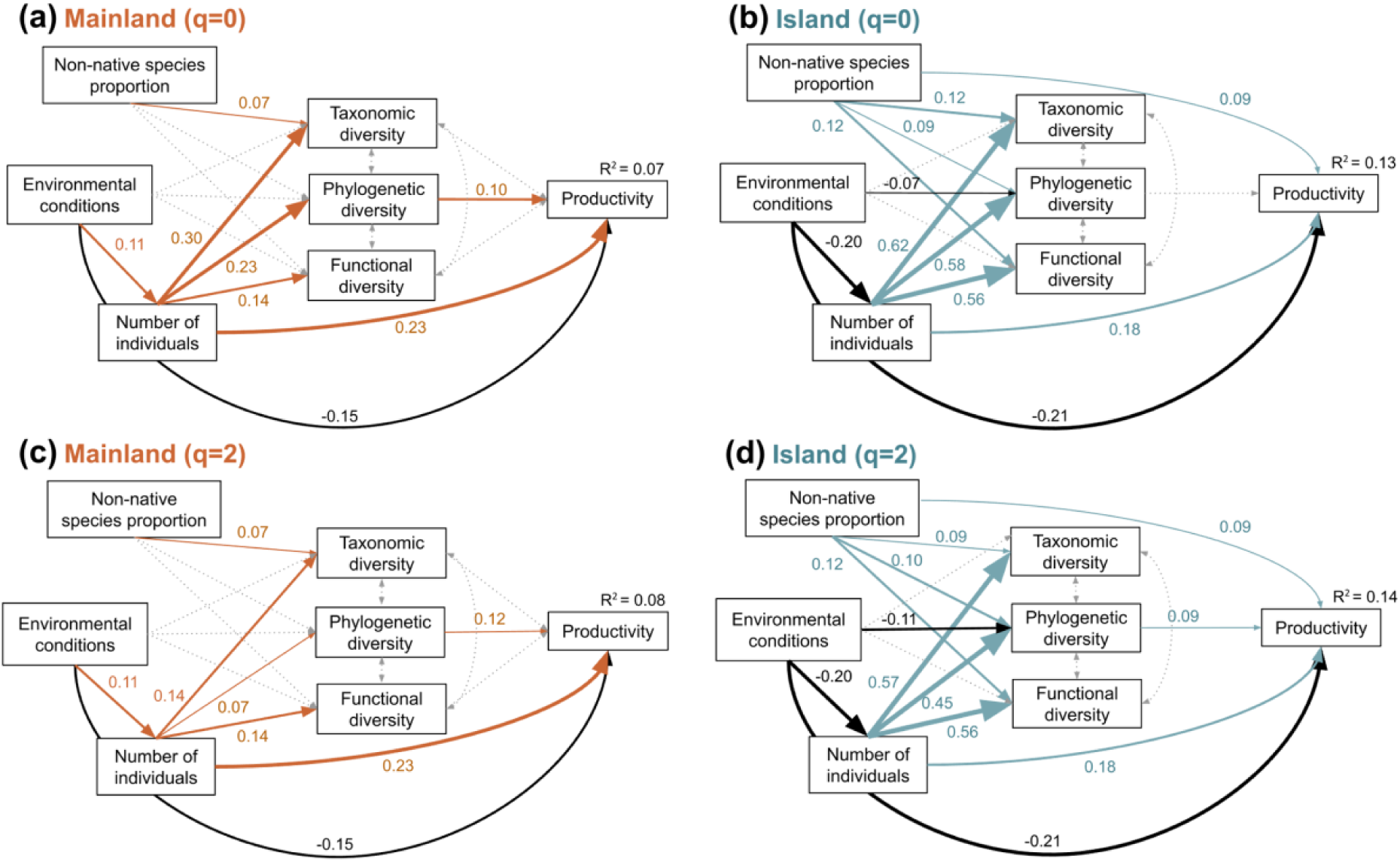
Piecewise structural equation models (SEMs) testing the effects of taxonomic, phylogenetic, and functional diversity in Spain in regions classified as “temperate, dry summer, warm summer” (Csb) under the Köppen-Geiger climate classification. Separate SEMs were fitted for Hill numbers 0 (i.e., species richness; a, b) and 2 (i.e., emphasizing common species influence; c, d) on productivity in mainland (a, c) and island (b, d) forests, when considering environmental conditions, proportion of non-native species, and number of individuals. Colored arrows indicate positive relationships, black arrows negative ones, dotted arrows show non-significant paths, and double-headed dotted arrows are correlations between variables. Standardized path coefficients are given next to significant paths. Line thickness of significant paths is scaled by the standardized path coefficients. (a) Fisher’s C = 4.104, df = 2, P-value = 0.13, and n = 1185; (b) Fisher’s C = 5.823, df = 4, P-value = 0.21, and n = 628; (c) Fisher’s C = 4.512, df = 2, P-value = 0.11, and n = 1185; and (d) Fisher’s C = 8.351, df = 4, P-value = 0.08, and n = 628.

All analyses were conducted in R version 4.2.1 (R Core Team, 2022). In addition to the already mentioned packages, we used the following packages: ‘dplyr’ (Wickham et al., 2020), ‘tidyr’ (Wickham & Henry, 2019), ‘stringr’ (Wickham, 2019), ‘raster’ (Hijmans, 2022), ‘FactoMineR’ (Le et al., 2008), ‘factoextra’ (Kassambara & Alboukadel, 2020), ‘performance’ (Lüdecke et al., 2021), ‘sp’ (Pebesma & Bivand, 2005), ‘spdep’ (Bivand, 2022), ‘kableExtra’ (Zhu, 2021), ‘ggplot2’ (Wickham, 2016), and ‘patchwork’ (Pedersen, 2020).

## Results

Annual volume increment, a proxy for productivity, was very similar for both island and mainland forests (Figure 3a). In contrast, overall taxonomic, phylogenetic, and functional diversity was higher for both Hill numbers (q=0 and q=2) in mainland forests compared with island forests (Figure 3b-g). Functional diversity displayed the greatest difference between island and mainland forests for both Hill numbers (q=0 and q=2; Figure 3f-g).

**Figure 3.**
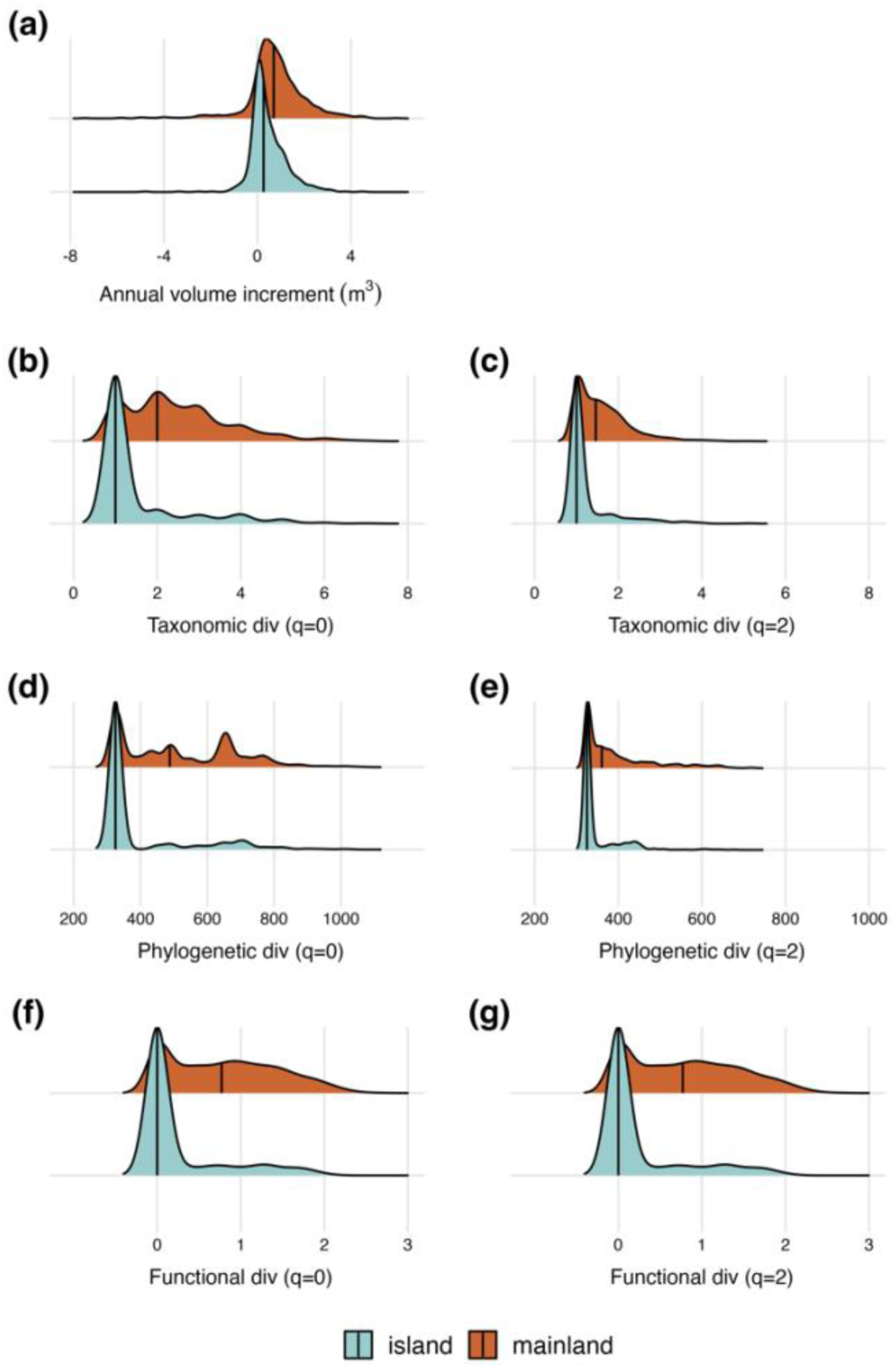
Distributions of plot-level productivity and multifaceted diversity indices in mainland and island forests in Spain in regions classified as “temperate, dry summer, warm summer” (Csb) under the Köppen-Geiger climate classification. Specifically, annual volume increment (m^3^) as a proxy for productivity (a), taxonomic diversity for Hill numbers 0 (b) and 2 (c), phylogenetic diversity for Hill numbers 0 (d) and 2 (e), and functional diversity for Hill numbers 0 (f) and 2 (g). Black line in each distribution indicates the median value.

When analyzing the effect of diversity facets on productivity, we found an overall significant positive effect of diversity on productivity (Figure 4; Table S4, p-value < 0.01) for all diversity facets for both Hill numbers (q=0 and q=2). The biogeographic context, i.e., mainland or island, had a significant effect on productivity only for taxonomic and phylogenetic diversity models when giving emphasis to common species (q=2; Table S4, p-value < 0.01), where mainland forests showed higher productivity values than islands. However, we did not find a significant interactive effect of biodiversity and biogeographic context.

Our SEMs showed that all models fitted the data well (p-value of the chi-square > 0.05; Tables S6-S9). Phylogenetic diversity had a positive effect on productivity in mainland forests (standardized path coefficients of direct effects = 0.10 and 0.12 for q=0 and 2, respectively; Figure 5a, c; total significant effect = 0.012 for q=1; Table S10). In contrast, phylogenetic diversity did not affect productivity (q=0; Figure 5b) or had a weak positive effect on island forest productivity when accounting for species abundances (q=2; standardized path coefficient range of direct effect = 0.09; Figure 5d).

On islands, the proportion of non-native species had a weak, yet positive direct effect on productivity (standardized path coefficient of direct effect = 0.09 for q = 0 and 2; Figures 5b, d), as well as via positive, indirect effects on phylogenetic diversity when giving emphasis to abundant species (standardized path coefficient of indirect effect = 0.008 for q = 2; Figure 5d). Environmental conditions influenced productivity via direct and indirect mechanisms across all the models (Figure 5; total significant effect = -0.12 for mainland for q=0 and 2, -0.25 for islands for q=0 and 2; Tables S10-S13). Specifically, productivity decreased in island forests under warmer and drier conditions either via negative effects on the number of individuals (standardized path coefficient of indirect effect = -0.21 for islands q=0 and 2; Figure 5b, d), phylogenetic diversity on islands when emphasizing abundant species (standardized path coefficient of indirect effect = -0.01; Figure 5d) or by alternative mechanisms in both island and mainland forests (standardized path coefficient of direct effect = -0.15 for mainland q=0 and 2, -0.21 for islands for q=0 and 2; Figure 5). While productivity increased under the same environmental conditions via positive effects on number of individuals in mainland forests (standardized path coefficient of indirect effect = 0.03 for mainland q=0 and 2; Figure 5a, c). In all models, the number of individuals showed positive direct effects on productivity (standardized path coefficient of direct effect = 0.23 for mainland q=0 and 2, 0.18 for islands q=0 and 2; Figure 5), and positive, indirect effects in mainland and island forests via phylogenetic diversity (standardized path coefficient of indirect effect = 0.05 and 0.02 for mainland q=0 and 2, for islands q=2; Figure 5a, c, d). Additionally, the number of individuals had the strongest total effect on productivity in mainland forests across all variables tested (total significant effect = 0.22 for q=0 and 2; Tables S10 and S12). While in island forests, environmental conditions had the overall strongest effects on productivity (total significant effect = -0.25 for q=0 and 2; Tables S11 and S13).

## Discussion

Our results reveal that the biogeographic context influences the magnitude of the biodiversity-productivity relationships, largely due to the direct and indirect impacts of non-native species on productivity. Furthermore, our results support the relevance of a multifaceted perspective when assessing BEF relationships in naturally-assembled ecosystems, through the influence of phylogenetic diversity on productivity that emerged as a result of variation in environmental conditions and non-native species. Our results provide evidence that a systematic understanding of BEF in naturally-assembled forests, regardless of the biogeographic context, requires the careful inclusion of other potentially confounding factors, such as environmental conditions and number of individuals.

We find that the biogeographic context contributes to the magnitude of BEF relationships, with the influence of the biogeographic context not being limited to environmental conditions, e.g., variation in temperature, precipitation, soil characteristics (Fanin et al., 2018; Ratcliffe et al., 2017). Specifically, our results show that in broadly similar climatic regions, mainland and island forests exhibit similar levels of productivity, yet island ecosystems display overall lower diversity values but stronger BEF relationships. This finding is in line with previous studies that have found comparatively lower species richness on islands than mainlands, a pattern explained mainly by island area and isolation at macroecological scales (Kreft et al., 2008; Whittaker & Fernández-Palacios, 2007) and by island age at local scales (Craven et al., 2019). Biogeographical factors could also influence the relative importance of underlying ecological processes, e.g., complementarity vs. selection effects, as certain native species may have disproportionate effects on ecosystem functioning, such as the highly abundant *P. canariensis* in the Canaries (Dee et al., 2023). Additionally, geographic isolation makes island ecosystems particularly susceptible to biological invasions (Sax et al., 2002) relative to mainland ecosystems, creating novel ecosystems that are usually more productive than non-invaded ones (Mascaro et al., 2012). Our results suggest that non-native species drive forest productivity in island forests via the phylogenetic novelty that they contribute to these novel ecosystems.

Biogeographical factors such as isolation cause islands to have empty niche space that is not filled by native species. Non-native species can fill these empty niches and promote biodiversity in island ecosystems (Sax & Gaines, 2008; Whittaker & Fernández-Palacios, 2007). Positive impacts of non-native species on ecosystem functioning are expected and could be attributed to their - potentially - novel traits or to enemy release, resulting from the absence of competitors that could suppress these non-native species populations. Where empty niches might be explained by evolutionary rates not being fast enough to fill these niches via speciation, or that the niches were once occupied by now extinct species, where the latter is less likely as plant invasions in islands have been found to trigger - unlike other taxa - almost no extinctions (Sax & Gaines, 2008). Furthermore, we found that non-native species on islands (e.g., *Pinus radiata*) not only affect productivity via phylogenetic diversity, but also directly. This pattern may suggest that recently introduced species are rarer and have a disproportionate impact due to the absence of natural enemies or competitors. This finding supports the enemy-release hypothesis, where species present for a longer time in islands may have accumulated natural enemies, leading to lower per capita impacts (see DeWalt et al., 2004).

Overall, biodiversity-productivity relationships were weak or even absent (e.g., phylogenetic diversity in island forests (q=0)) and only limited to phylogenetic diversity. These findings align with those of Craven et al. (2020), but contradict the ones of Ruiz-Benito et al. (2014), although they focused uniquely on taxonomic diversity and include wide climatic gradients. Yet, it is noteworthy that the spatial grain of plots may render smaller plots more vulnerable to the impact of demographic stochasticity, i.e., sampling variability in births and deaths (Storch & Okie, 2019). In addition, non-native species histories of introduction in mainland forests may differ from islands (van Kleunen et al., 2015), causing non-native species to have more time to accumulate competitors and, therefore, contribute less to productivity levels despite their contribution to diversity. In this regard, the biogeographic context of mainland forests may play a significant role in shaping the ecological dynamics of these ecosystems. Due to greater connectivity with other regions, niche space could be more evenly filled, thereby limiting the potential for non-native species to establish themselves. Non-native species showed a weak effect on taxonomic diversity in mainland forest, while their effects on island ecosystems were stronger and multifaceted. This suggests that non-native species in more diverse mainland forests provide less added diversity and less evolutionary or functional novelty than on (relatively species poor) islands, resulting in lower impact on ecosystem functioning.

The number of individuals consistently affected multifaceted diversity and productivity in mainland and island forests, which supports the more-individuals hypothesis. Our findings extend the commonly studied influence of the number of individuals on species richness, to phylogenetic and functional diversity. In both mainland and island forests, the number of individuals directly influenced productivity, and further regulated it by influencing phylogenetic diversity. This suggests that ecological processes, such as secondary succession - the process of recovery from natural or anthropogenic disturbances during which stem density increases non-linearly - are a fundamental determinant of ecosystem functioning for both island and mainland forests (Arroyo-Rodríguez et al., 2017; Chazdon et al., 2009; Gilroy et al., 2014). Further, our results highlight the potential benefits of increasing tree density on diversity and productivity, which may serve as a proxy for ecological processes that are important for restoring previously disturbed ecosystems. For instance, restoration efforts have the capacity to increase diversity and ecosystem functioning in degraded ecosystems (Benayas et al., 2009). Moreover, an increase in diversity through ecological restoration can reduce competition from invasive species and aid with the recovery of endangered native ecosystems such as island forests. On the other hand, environmental conditions have proven to moderately affect both forest diversity and productivity, acting as a limiting factor for both island and mainland ecosystems. Although the impacts of environmental conditions are primarily restricted to forest phylogenetic diversity on islands, this finding is consistent with the species-energy hypothesis linking climatic and soil resource availability to higher levels of diversity, which in turn enhances productivity in island forests. Our finding that environmental conditions directly influence forest productivity supports the findings of previous studies on the context-dependent nature of BEF relationships (Fanin et al., 2018; Guerrero-Ramírez et al., 2019; Ratcliffe et al., 2017; Tilman et al., 2014).

In conclusion, we show that biogeographic context mediates the biodiversity-productivity relationship in forests. Furthermore, we identified multiple pathways, i.e., phylogenetic diversity, the abundance of non-native species, environmental conditions, and number of individuals as universal drivers of the biodiversity-productivity relationship of forests in both geographical settings. However, we found the strength of these mechanisms to be different in mainland and island forests, suggesting that the relative importance of biogeographic context is greater than that of environmental conditions. The relevance of environmental conditions and biodiversity on ecosystem functioning in highly threatened ecosystems such as islands needs to be considered to better protect island forests from anthropogenic impacts and climate change.

## Acknowledgments

MLT acknowledges funding from the Studienstiftung des deutschen Volkes. The trait data collection on Tenerife was supported by the Deutsche Forschungsgemeinschaft (DFG) Research Training Group 1644 ‘Scaling Problems in Statistics’, grant no. 152112243 to HK and MPPB. NRG-R thanks the Dorothea Schlözer Postdoctoral Programme of the Georg-August-Universität Göttingen and the DFG, grant number 316045089/GRK 2300 for their support. AA is funded by a Serra-Hunter fellowship of the Generalitat de Catalunya. DC acknowledges funding from FONDECYT Project 1201347 and the Data Observatory Foundation.

## Data accessibility statement

All aggregated data and R codes used for analyses will become openly available upon publication.

## Notes

### Competing Interest Statement

The authors have declared no competing interest.

