## Supplementary information for "Biogeographic context mediates multifaceted diversity-productivity relationships in island and mainland forests"

**Running title:** Biodiversity-productivity relationships in islands and mainland forests

**Authors:** Maria Laura Tolmos[1], Nathaly R. Guerrero-Ramirez[1,2,3], Aitor Ameztegui[4,5], Martha Paola Barajas Barbosa[6,7], Dylan Craven[8,9,†], Holger Kreft[1,3,10†]

**Affiliations:**
[1] Biodiversity, Macroecology and Biogeography, University of Göttingen, Büsgenweg 1, 37077 Göttingen, Germany
[2] Silviculture and Forest Ecology of the Temperate Zones, University of Göttingen, Büsgenweg 1, 37077 Göttingen, Germany
[3] Centre for Biodiversity and Land Use (CBL), University of Göttingen, Büsgenweg 1, 37077 Göttingen, Germany
[4] Department of Agricultural and Forest Sciences and Engineering (DCEFA), University of Lleida, Av. Alcalde Rovira Roure 191, 25198 Lleida, Spain
[5] Joint Research Unit CTFC-AGROTECNIO-CERCA, Ctra. St. Llorenç de Morunys km 2, 25280 Solsona, Catalonia, Spain
[6] German Centre for Integrative Biodiversity Research (iDiv) Halle-Jena-Leipzig, Germany
[7] Department of Computer Science, Martin Luther University Halle-Wittenberg, Germany
[8] Centro de Modelación y Monitoreo de Ecosistemas, Universidad Mayor, José Toribio Medina 29, Santiago 8340589, Chile
[9] Data Observatory Foundation, Eliodoro Yáñez 2990, oficina A5, Providencia, Santiago, Chile
[10] Campus Institute Data Science (CIDAS), University of Göttingen, Goldschmidtstraße 1, 37077 Göttingen, Germany

**Corresponding author:** Maria Laura Tolmos; Biodiversity, Macroecology and Biogeography, University of Göttingen, Büsgenweg 1, 37077 Göttingen, Germany


† Joint senior authors

### Supplementary Information

**Table S1.** Tree species list for forest inventory plots in mainland and island forests in Spain in regions classified as "temperate, dry summer, warm summer" (Csb) under the Köppen-Geiger climate classification.

| Location | Species |
|---|---|
| Island | *Pinus canariensis* |
| Island | *Pinus pinea* |
| Island | *Pinus radiata* |
| Island | *Ulmus minor* |
| Island | *Prunus* |
| Island | *Salix pedicellata* |
| Island | *Castanea sativa* |
| Island | *Pinus halepensis* |
| Island | *Erica arborea* |
| Island | *Persea indica* |
| Island | *Myrica faya* |
| Island | *Laurus azorica* |
| Island | *Ilex canariensis* |
| Island | *Heberdenia bahamensis* |
| Island | *Eucalyptus globulus* |
| Island | *Ilex perado* |
| Island | *Prunus lusitanica* |
| Island | *Erica scoparia* |
| Island | *Picconia excelsa* |
| Island | *Arbutus canariensis* |
| Island | *Persea barbujana* |
| Island | *Arbutus unedo* |
| Island | *Juniperus turbinata* |
| Island | *Phoenix canariensis* |
| Island | *Rhamnus glandulosa* |
| Island | *Visnea mocanera* |
| Mainland | *Pinus pinaster* |
| Mainland | *Eucalyptus globulus* |
| Mainland | *Betula pubescens* |
| Mainland | *Pinus radiata* |
| Mainland | *Quercus robur* |
| Mainland | *Laurus nobilis* |
| Mainland | *Alnus glutinosa* |
| Mainland | *Salix atrocinerea* |
| Mainland | *Castanea sativa* |
| Mainland | *Ilex aquifolium* |
| Mainland | *Robinia pseudoacacia* |

| | |
|---|---|
| Mainland | *Pyrus* |
| Mainland | *Frangula alnus* |
| Mainland | *Betula* |
| Mainland | *Salix elaeagnos* |
| Mainland | *Salix* |
| Mainland | *Quercus pyrenaica* |
| Mainland | *Quercus petraea* |
| Mainland | *Populus nigra* |
| Mainland | *Eucalyptus gomphocephala* |
| Mainland | *Salix caprea* |
| Mainland | *Pinus sylvestris* |
| Mainland | *Cedrus deodara* |
| Mainland | *Quercus suber* |
| Mainland | *Juglans regia* |
| Mainland | *Prunus* |
| Mainland | *Fraxinus angustifolia* |
| Mainland | *Eucalyptus camaldulensis* |
| Mainland | *Sambucus nigra* |
| Mainland | *Crataegus monogyna* |
| Mainland | *Acer pseudoplatanus* |
| Mainland | *Corylus avellana* |
| Mainland | *Fraxinus excelsior* |
| Mainland | *Crataegus* |
| Mainland | *Arbutus unedo* |
| Mainland | *Cornus sanguinea* |
| Mainland | *Quercus ilex* |
| Mainland | *Sorbus aucuparia* |
| Mainland | *Populus x canadensis* |
| Mainland | *Acacia dealbata* |
| Mainland | *Pinus pinea* |
| Mainland | *Ulmus glabra* |
| Mainland | *Ulmus minor* |
| Mainland | *Cupressus sempervirens* |
| Mainland | *Acacia melanoxylon* |
| Mainland | *Sorbus* |
| Mainland | *Malus sylvestris* |
| Mainland | *Eucalyptus viminalis* |
| Mainland | *Eucalyptus nitens* |
| Mainland | *Acer opalus* |
| Mainland | *Quercus pubescens* |
| Mainland | *Fagus sylvatica* |
| Mainland | *Sorbus aria* |
| Mainland | *Acer monspessulanum* |
| Mainland | *Acer campestre* |
| Mainland | *Platanus hispanica* |

| | |
|---|---|
| Mainland | *Pinus halepensis* |
| Mainland | *Quercus faginea* |
| Mainland | *Pinus nigra* |
| Mainland | *Populus alba* |
| Mainland | *Phillyrea latifolia* |
| Mainland | *Ficus carica* |
| Mainland | *Salix euxina* |
| Mainland | *Juniperus oxycedrus* |
| Mainland | *Taxus baccata* |
| Mainland | *Juniperus phoenicea* |
| Mainland | *Salix alba* |
| Mainland | *Eucalyptus robusta* |
| Mainland | *Prunus avium* |
| Mainland | *Pseudotsuga menziesii* |
| Mainland | *Acacia* |
| Mainland | *Populus tremula* |

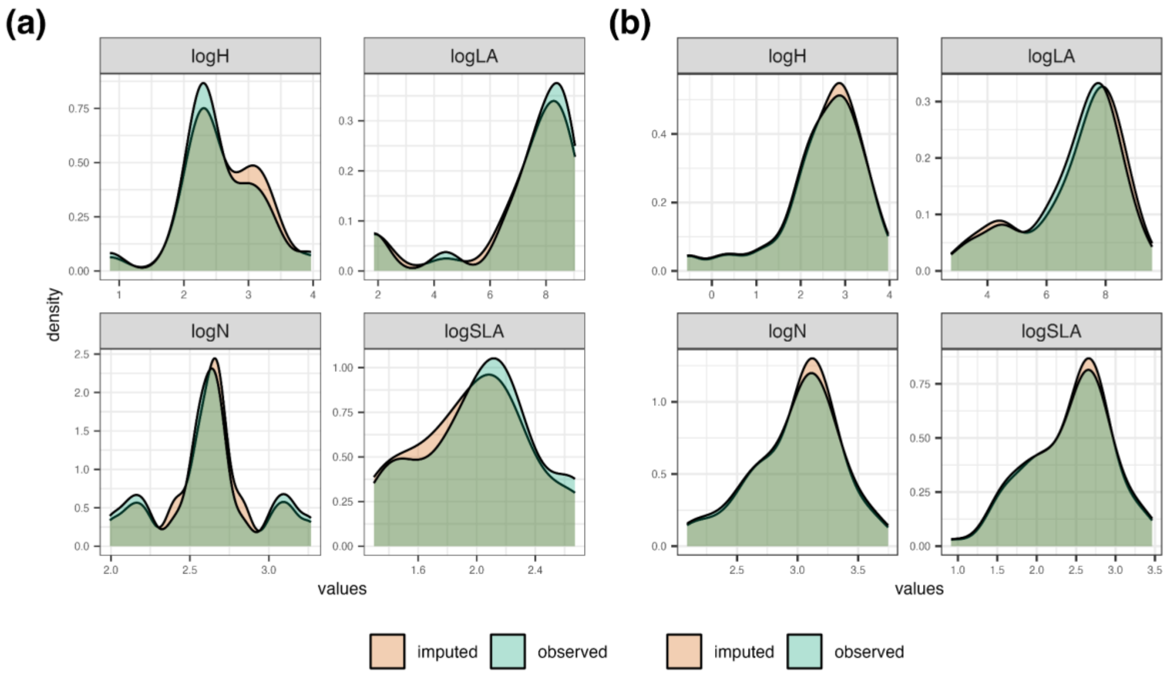


**Figure S1.** Observed and imputed traits distributions on mainland (a) and island (b) forests in Spain in regions classified as "temperate, dry summer, warm summer" (Csb) under the Köppen-Geiger climate classification. The four functional traits chosen reflect plant acquisitive strategies (Díaz et al., 2016; Reich, 2014; Wright et al., 2004): maximum height (H; m), specific leaf area (SLA; $mm^2$/mg), leaf area (LA; $mm^2$), and leaf nitrogen content per leaf dry mass ($N_{mass}$; mg/g).

**Table S2.** Mean squared errors (MSE) for each trait using a phylogenetic imputation approach in **mainland forests**: maximum height (H), specific leaf area (SLA), leaf area (LA), and leaf nitrogen content per leaf dry mass ($N_{mass}$), and phylogenetic eigenvectors (k). Bold k and MSE values correspond to the lowest error for each trait. Please note that imputation error is in the unit of each functional trait. Bottom line section in gray shows the normalized root mean square error (NRMSE) for each trait following Penone et al. (2014).

| k | H error | LA error | $N_{mass}$ error | SLA error |
|---|---|---|---|---|
| 1 | 177.56 | 6217373.28 | 31.26 | 21.53 |
| **2** | 160.21 | 6723532.34 | 26.13 | **20.27** |
| 3 | 139.59 | 6306972.78 | 23.53 | 22.31 |
| 4 | 129.27 | 6563767.34 | 24.32 | 21.55 |
| 5 | 134.30 | 6660207.15 | 25.70 | 26.08 |
| 6 | 134.52 | 6458549.10 | 21.13 | 23.17 |
| 7 | 129.03 | 5860089.09 | 23.33 | 23.49 |
| 8 | 131.03 | 6277377.82 | 20.67 | 23.84 |
| 9 | 130.68 | 5799856.91 | 20.84 | 25.11 |
| **10** | 131.26 | 6118571.42 | **19.92** | 23.56 |
| 11 | 124.23 | 6298539.01 | 20.18 | 23.79 |
| 12 | 119.90 | 6078260.34 | 20.23 | 26.05 |
| 13 | 119.60 | 5925402.67 | 21.59 | 23.81 |
| 14 | 121.09 | 6047135.36 | 23.30 | 26.23 |
| 15 | 116.58 | 5974085.55 | 21.58 | 27.40 |
| 16 | 116.06 | 5631866.67 | 20.20 | 26.43 |
| 17 | 112.03 | 6127369.21 | 21.61 | 27.73 |
| 18 | 112.57 | 5813140.92 | 23.54 | 26.61 |
| 19 | 102.00 | 6421662.58 | 21.27 | 26.95 |
| 20 | 102.54 | 5776962.88 | 24.11 | 27.18 |
| 21 | 108.11 | 5783277.88 | 23.54 | 26.62 |
| 22 | 114.24 | 6334204.83 | 21.88 | 25.80 |
| **23** | **99.99** | 5728846.86 | 21.14 | 27.66 |
| 24 | 108.21 | 6302132.95 | 22.73 | 27.54 |
| 25 | 105.93 | 5917915.67 | 22.44 | 26.75 |
| 26 | 115.88 | 6643944.88 | 24.24 | 28.49 |
| 27 | 109.03 | 5906109.64 | 24.72 | 27.37 |
| 28 | 116.70 | 5942372.54 | 21.46 | 27.30 |
| 29 | 110.13 | 6033371.82 | 23.72 | 26.91 |
| 30 | 104.50 | 5936010.32 | 24.27 | 29.19 |
| 31 | 109.29 | 6052335.73 | 22.83 | 27.24 |
| 32 | 106.92 | 6522725.61 | 22.31 | 28.32 |
| 33 | 121.80 | 6554456.98 | 24.45 | 31.23 |
| 34 | 108.61 | 5965649.01 | 23.36 | 26.50 |
| 35 | 122.00 | 5707243.47 | 23.32 | 27.80 |
| 36 | 108.14 | 5660194.35 | 25.06 | 26.85 |

| | | | | |
|---|---|---|---|---|
| 37 | 126.40 | 5647690.39 | 22.57 | 30.78 |
| 38 | 122.22 | 5622682.02 | 24.83 | 27.79 |
| 39 | 114.88 | 5958053.67 | 24.94 | 27.18 |
| 40 | 124.21 | 6030135.54 | 22.57 | 27.66 |
| 41 | 120.87 | 5492868.67 | 22.43 | 27.73 |
| 42 | 118.84 | 5819531.29 | 23.51 | 26.70 |
| 43 | 130.22 | 6138485.86 | 22.03 | 27.88 |
| 44 | 125.70 | 5397437.37 | 23.44 | 27.85 |
| 45 | 119.86 | 5285033.12 | 25.39 | 27.82 |
| 46 | 120.88 | 5851395.96 | 23.69 | 28.06 |
| 47 | 125.73 | 5783681.71 | 25.35 | 27.56 |
| 48 | 115.62 | 5138717.90 | 22.81 | 26.81 |
| 49 | 127.37 | 5226885.23 | 24.98 | 30.02 |
| 50 | 122.01 | 5113427.13 | 23.83 | 27.84 |
| **51** | 119.26 | **4539018.32** | 24.18 | 27.21 |
| 52 | 121.58 | 4856048.56 | 23.19 | 26.67 |
| 53 | 117.64 | 5568216.67 | 23.74 | 26.04 |
| 54 | 120.46 | 5147050.44 | 25.82 | 27.08 |
| 55 | 125.71 | 4578190.09 | 25.58 | 27.49 |
| 56 | 119.24 | 4902928.85 | 24.17 | 26.65 |
| 57 | 118.08 | 5271631.26 | 26.55 | 26.41 |
| 58 | 118.04 | 5075072.06 | 27.16 | 26.66 |
| 59 | 113.72 | 5521209.80 | 26.90 | 25.86 |
| 60 | 120.89 | 5074658.50 | 24.76 | 27.98 |
| 61 | 126.90 | 5284274.54 | 23.34 | 26.55 |
| 62 | 126.71 | 5061790.13 | 26.00 | 25.16 |
| 63 | 124.48 | 4865250.95 | 24.36 | 24.68 |
| 64 | 129.39 | 5758627.73 | 26.25 | 24.09 |
| 65 | 121.98 | 5247837.17 | 24.57 | 22.88 |
| 66 | 137.83 | 6040974.16 | 26.73 | 25.14 |
| 67 | 123.68 | 5504621.78 | 23.55 | 27.00 |
| 68 | 119.20 | 5287389.13 | 25.33 | 26.59 |
| 69 | 132.84 | 5652540.97 | 26.49 | 25.21 |
| **NRMSE** | 1.38 | 17.78 | 0.76 | 0.83 |

**Table S3.** Mean squared errors (MSE) for each trait using a phylogenetic imputation approach in **island forests**: maximum height (H), specific leaf area (SLA), leaf area (LA), and leaf nitrogen content per leaf dry mass ($N_{mass}$) , and phylogenetic eigenvectors (k). Bold k and MSE values correspond to the lowest error for each trait. Please note that imputation error is in the unit of each functional trait. Bottom line section in gray shows the normalized root mean square error (NRMSE) for each trait following Penone et al. (2014).

| **k** | **H error** | **LA error** | **$N_{mass}$ error** | **SLA error** |
|---|---|---|---|---|
| 1 | 124.91 | 8228871.35 | 20.18 | 10.04 |
| 2 | 124.06 | 7887372.32 | 16.63 | 10.95 |
| 3 | 120.31 | 7407288.80 | 20.82 | 12.21 |
| 4 | 139.63 | 6583222.34 | 17.23 | 10.84 |
| 5 | 137.57 | 7581605.15 | 18.12 | 9.86 |
| 6 | 145.39 | 6876115.60 | 18.44 | 9.44 |
| 7 | 148.90 | 6694910.81 | 17.43 | 10.46 |
| 8 | 130.31 | 6567551.86 | 18.38 | 9.22 |
| 9 | 147.71 | 6733981.75 | 19.49 | 11.28 |
| 10 | 136.77 | 6897188.23 | 18.63 | 10.88 |
| **11** | 153.39 | **6338468.22** | 18.94 | 9.09 |
| **12** | 149.11 | 7462710.77 | **16.42** | 10.43 |
| 13 | 123.79 | 7058994.81 | 18.21 | 10.67 |
| 14 | 123.29 | 7585807.34 | 18.78 | 11.16 |
| 15 | 114.29 | 8074766.08 | 20.86 | 10.46 |
| 16 | 102.17 | 7808207.94 | 21.87 | 11.69 |
| 17 | 116.01 | 6911854.53 | 21.03 | 11.80 |
| 18 | 118.35 | 7174041.22 | 22.11 | 11.82 |
| **19** | **96.45** | 6344967.62 | 19.35 | 10.70 |
| 20 | 112.46 | 6935631.52 | 20.81 | 11.03 |
| 21 | 114.35 | 6834221.97 | 21.40 | 11.43 |
| 22 | 99.17 | 7088914.86 | 21.76 | 10.77 |
| 23 | 113.96 | 7388479.51 | 21.51 | 9.51 |
| **24** | 102.25 | 7285148.52 | 18.42 | **8.82** |
| **NRMSE** | 1.38 | 26.96 | 0.97 | 0.93 |

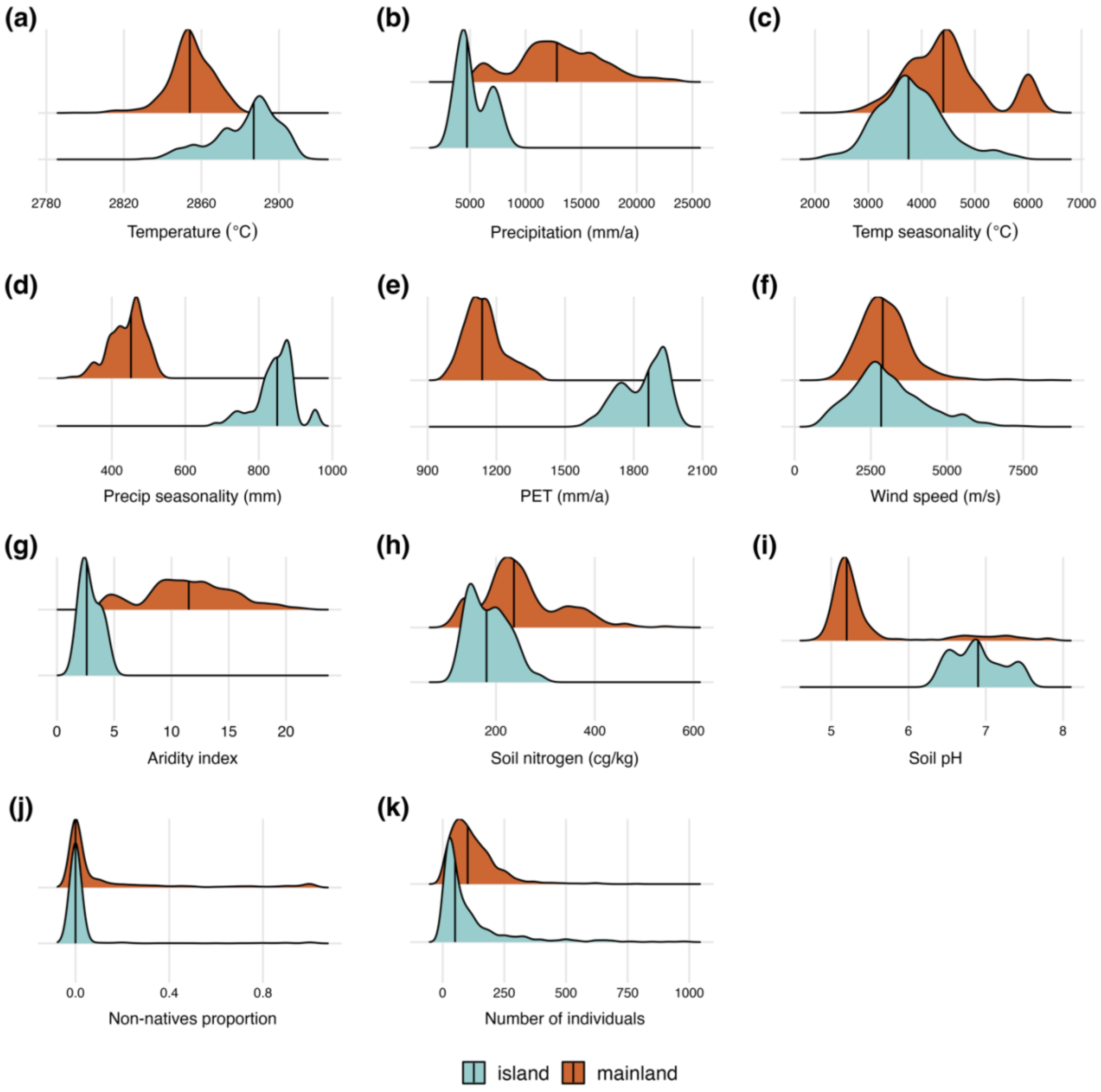


**Figure S2.** Distributions of variables in mainland and island forests in Spain in regions classified as "temperate, dry summer, warm summer" (Csb) under the Köppen-Geiger climate classification. Specifically, environmental conditions (a-i), non-native species proportion per plot (j), and number of individuals per plot (k). Black line in each distribution indicates the median value.

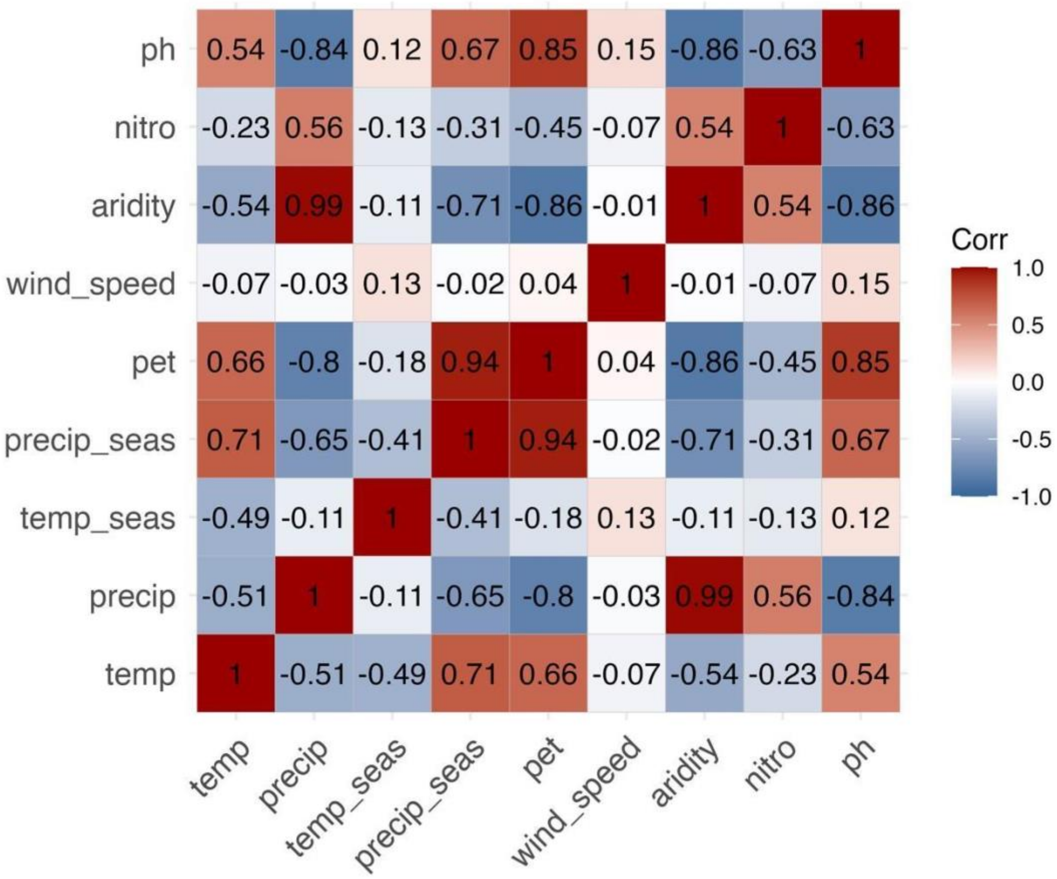


**Figure S3.** Pairwise correlation matrix for environmental variables in mainland and island forests in Spain in regions classified as “temperate, dry summer, warm summer” (Csb) under the Köppen-Geiger climate classification.

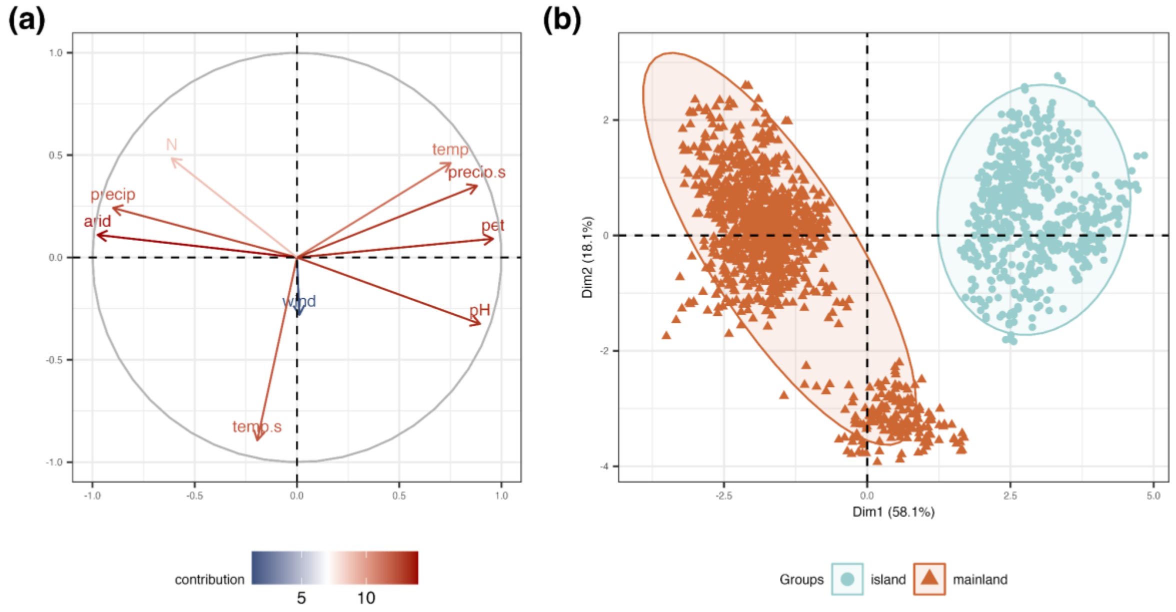


**Figure S4.** Principal component analysis (PCA) of environmental conditions influencing tree diversity and productivity in mainland and island forests in Spain in regions classified as "temperate, dry summer, warm summer" (Csb) under the Köppen-Geiger climate classification. (a) Environmental conditions contributions where red shows a higher contribution and blue a lower one. (b) Triangles and dots represent plots for mainland (orange triangles) and island (blue dots) forests, distributed across the first two axes of the PCA.

**Table S4.** Summary of linear models examining the influence of taxonomic, phylogenetic, and functional diversity on productivity for Hill numbers 0 and 2 in mainland and island forests in Spain in regions classified as "temperate, dry summer, warm summer" (Csb) under the Köppen-Geiger climate classification.

| **Taxonomic diversity (q=0)** | | | | | | **Taxonomic diversity (q=2)** | | | | | |
|---|---|---|---|---|---|---|---|---|---|---|---|
| | Estimate | Sum Sq | Df | F value | p-value | | Estimat e | Sum Sq | Df | F value | p-value |
| (Intercept) | -0.08 | 2.60 | 1 | 2.69 | 0.10 | (Intercept) | -0.12 | 6.82 | 1 | 7.00 | 0.01 |
| tax div0 | 0.17 | 14.28 | 1 | 14.75 | 0.00 | tax div2 | 0.15 | 10.92 | 1 | 11.22 | 0.00 |
| location | 0.14 | 5.78 | 1 | 5.97 | 0.01 | location | 0.19 | 12.94 | 1 | 13.30 | 0.00 |
| tax div0:location | -0.06 | 1.03 | 1 | 1.06 | 0.30 | tax div2:location | -0.06 | 1.29 | 1 | 1.33 | 0.25 |
| Residuals | | 1751.98 | 1809 | | | Residuals | | 1760.26 | 1809 | | |
| **Phylogenetic diversity (q=0)** | | | | | | **Phylogenetic diversity (q=2)** | | | | | |
| | Estimate | Sum Sq | Df | F value | p-value | | Estimat e | Sum Sq | Df | F value | p-value |
| (Intercept) | -0.09 | 3.59 | 1 | 3.73 | 0.05 | (Intercept) | -0.09 | 3.66 | 1 | 3.79 | 0.05 |
| phylo div0 | 0.19 | 16.83 | 1 | 17.47 | 0.00 | phylo div2 | 0.23 | 16.26 | 1 | 16.85 | 0.00 |
| location | 0.15 | 7.44 | 1 | 7.72 | 0.01 | location | 0.16 | 8.71 | 1 | 9.02 | 0.00 |
| phylo div0:location | -0.06 | 1.22 | 1 | 1.27 | 0.26 | phylo div2:location | -0.12 | 3.71 | 1 | 3.84 | 0.05 |
| Residuals | | 1743.31 | 1809 | | | Residuals | | 1746.32 | 1809 | | |
| **Functional diversity (q=0)** | | | | | | **Functional diversity (q=2)** | | | | | |
| | Estimate | Sum Sq | Df | F value | p-value | | Estimat e | Sum Sq | Df | F value | p-value |
| (Intercept) | -0.08 | 2.96 | 1 | 3.05 | 0.08 | (Intercept) | -0.08 | 2.96 | 1 | 3.05 | 0.08 |
| func div0 | 0.19 | 13.26 | 1 | 13.69 | 0.00 | func div2 | 0.19 | 13.26 | 1 | 13.69 | 0.00 |
| location | 0.15 | 6.87 | 1 | 7.09 | 0.01 | location | 0.15 | 6.87 | 1 | 7.09 | 0.01 |
| func div0:location | -0.08 | 1.70 | 1 | 1.76 | 0.18 | func div2:location | -0.08 | 1.70 | 1 | 1.76 | 0.18 |
| Residuals | | 1752.39 | 1809 | | | Residuals | | 1752.39 | 1809 | | |

**Table S5.** Principal component analysis (PCA) of environmental conditions influencing tree diversity and productivity in mainland and island forests in Spain in regions classified as "temperate, dry summer, warm summer" (Csb) under the Köppen-Geiger climate classification. Eigenvalues and trait loadings of principal components (PC1 and PC2) in two different PCAs.

| | **PC1** | **PC2** |
|---|---|---|
| Variation explained (%) | 57.87 | 18.26 |
| Eigenvalue | 5.21 | 1.64 |
| **Variable loadings** | | |
| Temperature | 0.74 | 0.48 |
| Precipitation | -0.90 | 0.24 |
| PET | 0.96 | 0.09 |
| Wind velocity | 0.01 | -0.27 |
| Temperature seasonality | -0.17 | -0.90 |
| Precipitation seasonality | 0.88 | 0.35 |
| Aridity | -0.98 | 0.11 |
| Soil nitrogen | -0.62 | 0.48 |
| Soil pH | 0.90 | -0.31 |
| **Contributions (%)** | | |
| Temperature | 10.49 | 14.09 |
| Precipitation | 15.49 | 3.63 |
| PET | 17.63 | 0.53 |
| Wind velocity | 0.00 | 4.31 |
| Temperature seasonality | 0.55 | 49.15 |
| Precipitation seasonality | 14.81 | 7.50 |
| Aridity | 18.26 | 0.72 |
| Soil nitrogen | 7.31 | 14.11 |
| Soil pH | 15.44 | 5.95 |

**Table S6.** Standardized estimates of a structural equation models (SEM) exploring the direct and indirect influence of multifaceted diversity not considering species abundances (q=0), environmental conditions, number of individuals, and the proportion of non-native species on productivity in **mainland** forests in Spain in regions classified as "temperate, dry summer, warm summer" (Csb) under the Köppen-Geiger climate classification. P-values in bold letters are significant.

| **Mainland (q=0)** | | | | |
|---|---|---|---|---|
| **Response** | **Predictor** | **Std. Estimate** | **Standard error** | **p-value** |
| productivity | taxonomic div | -0.10 | 0.06 | 0.06 |
| $R^2$=0.07 | phylogenetic div | 0.10 | 0.11 | **0.04** |
| | functional div | 0.06 | 0.07 | 0.13 |
| | environmental conditions | -0.15 | -0.38 | **0.00** |
| | number of individuals | 0.23 | 0.30 | **0.00** |
| taxonomic div | environmental conditions | -0.04 | -0.09 | 0.12 |
| $R^2$=0.09 | number of individuals | 0.30 | 0.32 | **0.00** |
| | non-native species | 0.07 | 0.05 | **0.02** |
| phylogenetic div | environmental conditions | 0.00 | 0.00 | 0.99 |
| $R^2$=0.06 | number of individuals | 0.23 | 0.26 | **0.00** |
| | non-native species | -0.04 | -0.03 | 0.18 |
| functional div | environmental conditions | -0.05 | -0.12 | 0.08 |
| $R^2$=0.02 | number of individuals | 0.14 | 0.16 | **0.00** |
| | non-native species | 0.05 | 0.05 | 0.07 |
| number of individuals | environmental conditions | 0.11 | 0.22 | **0.00** |
| $R^2$=0.01 | | | | |
| **Partial correlations** | | | | |
| taxonomic div ~ phylogenetic div | | 0.78 | | **0.00** |
| taxonomic div ~ functional div | | 0.71 | | **0.00** |
| phylogenetic div ~ functional div | | 0.67 | | **0.00** |
| non-native species ~ number of individuals | | -0.13 | | **0.00** |
| Fisher's C = 4.104 | df = 2 | p-value = 0.13 | n = 1185 | |

**Table S7.** Standardized estimates of a structural equation models (SEM) exploring the direct and indirect influence of multifaceted diversity not considering species abundances (q=0), environmental conditions, number of individuals, and the proportion of non-native species on productivity in **island** forests in Spain in regions classified as "temperate, dry summer, warm summer" (Csb) under the Köppen-Geiger climate classification. P-values in bold letters are significant.

| **Island (q =0 )** | | | | |
|---|---|---|---|---|
| **Response** | **Predictor** | **Std. Estimate** | **Standard error** | **p-value** |
| productivity | phylogenetic div | 0.08 | 0.04 | 0.09 |
| $R^2$=0.13 | environmental conditions | -0.21 | 0.09 | **0.00** |
| | number of individuals | 0.18 | 0.03 | **0.00** |
| | non-native species | 0.09 | 0.04 | **0.01** |
| taxonomic div | environmental conditions | -0.03 | 0.09 | 0.28 |
| $R^2$=0.41 | number of individuals | 0.62 | 0.02 | **0.00** |
| | non-native species | 0.12 | 0.05 | **0.00** |
| phylogenetic div | environmental conditions | -0.07 | 0.09 | **0.03** |
| $R^2$=0.37 | number of individuals | 0.58 | 0.02 | **0.00** |
| | non-native species | 0.09 | 0.05 | **0.00** |
| functional div | environmental conditions | -0.05 | 0.09 | 0.13 |
| $R^2$=0.35 | number of individuals | 0.56 | 0.02 | **0.00** |
| | non-native species | 0.12 | 0.04 | **0.00** |
| number of individuals | environmental conditions | -0.20 | 0.15 | **0.00** |
| $R^2$=0.04 | | | | |
| **Partial correlations** | | | | |
| taxonomic div ~ phylogenetic div | | 0.92 | | **0.00** |
| taxonomic div ~ functional div | | 0.87 | | **0.00** |
| phylogenetic div ~ functional div | | 0.82 | | **0.00** |
| non-native species ~ number of individuals | | 0.05 | | 0.12 |
| Fisher's C = 5.823 | df = 4 | p-value = 0.21 | n = 628 | |

**Table S8.** Standardized estimates of a structural equation models (SEM) exploring the direct and indirect influence of multifaceted diversity accounting for species abundances (q=2), environmental conditions, number of individuals, and the proportion of non-native species on productivity in **mainland** forests in Spain in regions classified as "temperate, dry summer, warm summer" (Csb) under the Köppen-Geiger climate classification. P-values in bold letters are significant.

| MAINLAND (q=2) | | | | |
|---|---|---|---|---|
| **Response** | **Predictor** | **Std. Estimate** | **Standard error** | **p-value** |
| productivity | taxonomic div | -0.03 | 0.06 | 0.58 |
| $R^2$=0.08 | phylogenetic div | 0.12 | 0.05 | **0.01** |
| | functional div | -0.01 | 0.08 | 0.88 |
| | environmental conditions | -0.15 | 0.08 | **0.00** |
| | number of individuals | 0.23 | 0.04 | **0.00** |
| taxonomic div | environmental conditions | -0.03 | 0.07 | 0.30 |
| $R^2$=0.02 | number of individuals | 0.14 | 0.03 | **0.00** |
| | non-native species | 0.07 | 0.03 | **0.03** |
| phylogenetic div | environmental conditions | 0.05 | 0.07 | 0.10 |
| $R^2$=0.01 | number of individuals | 0.07 | 0.04 | **0.02** |
| | non-native species | -0.06 | 0.03 | 0.05 |
| functional div | environmental conditions | -0.05 | 0.07 | 0.08 |
| $R^2$=0.02 | number of individuals | 0.14 | 0.03 | **0.00** |
| | non-native species | 0.05 | 0.03 | 0.07 |
| number of individuals | environmental conditions | 0.11 | 0.06 | **0.00** |
| $R^2$=0.02 | | | | |
| **Partial correlations** | | | | |
| taxonomic div ~ phylogenetic div | | 0.64 | | **0.00** |
| taxonomic div ~ functional div | | 0.85 | | **0.00** |
| phylogenetic div ~ functional div | | 0.78 | | **0.00** |
| non-native species ~ number of individuals | | -0.13 | | **0.00** |
| Fisher's C = 4.512 | df = 2 | p-value = 0.11 | n = 1185 | |

**Table S9.** Standardized estimates of a structural equation models (SEM) exploring the direct and indirect influence of multifaceted diversity accounting for species abundances (q=2), environmental conditions, number of individuals, and the proportion of non-native species on productivity in **island** forests in Spain in regions classified as "temperate, dry summer, warm summer" (Csb) under the Köppen-Geiger climate classification. P-values in bold letters are significant.

| ISLAND (q=2) | | | | |
|---|---|---|---|---|
| **Response** | **Predictor** | **Std. Estimate** | **Standard error** | **p-value** |
| productivity | phylogenetic div | 0.09 | 0.04 | **0.03** |
| $R^2$=0.14 | environmental conditions | -0.21 | 0.09 | **0.00** |
| | number of individuals | 0.18 | 0.03 | **0.00** |
| | non-native species | 0.09 | 0.04 | **0.02** |
| taxonomic div | environmental conditions | -0.02 | 0.10 | 0.48 |
| $R^2$=0.35 | number of individuals | 0.57 | 0.03 | **0.00** |
| | non-native species | 0.09 | 0.05 | **0.00** |
| phylogenetic div | environmental conditions | -0.11 | 0.08 | **0.00** |
| $R^2$=0.25 | number of individuals | 0.45 | 0.02 | **0.00** |
| | non-native species | 0.10 | 0.04 | **0.00** |
| functional div | environmental conditions | -0.05 | 0.09 | 0.13 |
| $R^2$=0.35 | number of individuals | 0.56 | 0.02 | **0.00** |
| | non-native species | 0.12 | 0.04 | **0.00** |
| number of individuals | environmental conditions | -0.20 | 0.15 | **0.00** |
| $R^2$=0.04 | | | | |
| **Partial correlations** | | | | |
| taxonomic div ~ phylogenetic div | | 0.76 | | **0.00** |
| taxonomic div ~ functional div | | 0.93 | | **0.00** |
| phylogenetic div ~ functional div | | 0.83 | | **0.00** |
| non-native species ~ number of individuals | | 0.05 | | 0.12 |
| Fisher's C = 8.351 | df = 4 | p-value = 0.08 | n = 628 | |

**Table S10.** Direct, indirect, total, and mediator effects of a structural equation model (SEM) exploring the influence of multifaceted diversity not considering species abundances (q=0), environmental conditions, number of individuals (N), and the proportion of non-native species on productivity in **mainland** forests in Spain in regions classified as "temperate, dry summer, warm summer" (Csb) under the Köppen-Geiger climate classification. (*) shows significant effects.

| MAINLAND (q=0) | | | | | | | |
|---|---|---|---|---|---|---|---|
| | | **Effect** | **Bias** | **Std. Err.** | **Lower CI** | **Upper CI** | |
| DIRECT | environmental conditions | -0.144 | 0.000 | 0.023 | -0.190 | -0.098 | * |
| | taxonomic div | -0.053 | 0.000 | 0.027 | -0.105 | 0.001 | |
| | phylogenetic div | 0.058 | 0.000 | 0.028 | 0.003 | 0.111 | * |
| | functional div | 0.043 | 0.000 | 0.029 | -0.013 | 0.101 | |
| | number of individuals | 0.214 | 0.000 | 0.026 | 0.162 | 0.266 | * |
| INDIRECT | environmental conditions | 0.024 | 0.000 | 0.007 | 0.011 | 0.040 | * |
| | non-native species | -0.003 | 0.000 | 0.003 | -0.010 | 0.003 | |
| | number of individuals | 0.004 | 0.000 | 0.005 | -0.006 | 0.014 | |
| TOTAL | environmental conditions | -0.120 | 0.000 | 0.022 | -0.163 | -0.076 | * |
| | non-native species | -0.003 | 0.000 | 0.003 | -0.010 | 0.003 | |
| | taxonomic div | -0.053 | 0.000 | 0.027 | -0.105 | 0.001 | |
| | phylogenetic div | 0.058 | 0.000 | 0.028 | 0.003 | 0.111 | * |
| | functional div | 0.043 | 0.000 | 0.029 | -0.013 | 0.101 | |
| | number of individuals | 0.217 | 0.000 | 0.025 | 0.168 | 0.266 | * |
| MEDIATORS | taxonomic div | -0.018 | 0.000 | 0.010 | -0.039 | -0.001 | * |
| | phylogenetic div | 0.012 | 0.000 | 0.007 | 0.001 | 0.028 | * |
| | functional div | 0.007 | 0.000 | 0.005 | 0.000 | 0.022 | |
| | number of individuals | 0.024 | 0.000 | 0.007 | 0.012 | 0.039 | * |

**Table S11.** Direct, indirect, total, and mediator effects of a structural equation model (SEM) exploring the influence of multifaceted diversity not considering species abundances (q=0), environmental conditions, number of individuals (N), and the proportion of non-native species on productivity in **island** forests in Spain in regions classified as "temperate, dry summer, warm summer" (Csb) under the Köppen-Geiger climate classification. (*) shows significant effects.

| ISLAND (q=0) | | | | | | | |
|---|---|---|---|---|---|---|---|
| | | **Effect** | **Bias** | **Std. Err.** | **Lower CI** | **Upper CI** | |
| DIRECT | environmental conditions | -0.206 | 0.000 | 0.032 | -0.267 | -0.141 | * |
| | non-native species | 0.091 | 0.000 | 0.071 | -0.054 | 0.224 | |
| | phylogenetic div | 0.063 | 0.000 | 0.043 | -0.018 | 0.147 | |
| | number of individuals | 0.141 | 0.000 | 0.033 | 0.076 | 0.204 | * |
| INDIRECT | environmental conditions | -0.039 | 0.000 | 0.008 | -0.058 | -0.024 | * |
| | non-native species | 0.006 | 0.000 | 0.005 | -0.001 | 0.021 | |
| | number of individuals | 0.036 | 0.000 | 0.024 | -0.010 | 0.084 | |
| TOTAL | environmental conditions | -0.246 | 0.000 | 0.032 | -0.306 | -0.181 | * |
| | non-native species | 0.097 | 0.000 | 0.070 | -0.046 | 0.228 | |
| | phylogenetic div | 0.063 | 0.000 | 0.043 | -0.018 | 0.147 | |
| | number of individuals | 0.177 | -0.001 | 0.027 | 0.124 | 0.231 | |
| MEDIATORS | phylogenetic div | 0.030 | 0.000 | 0.021 | -0.008 | 0.074 | |
| | number of individuals | -0.035 | 0.000 | 0.008 | -0.052 | -0.021 | * |

**Table S12.** Direct, indirect, total, and mediator effects of a structural equation model (SEM) exploring the influence of multifaceted diversity accounting for species abundances (q=2), environmental conditions, number of individuals (N), and the proportion of non-native species on productivity in **mainland** forests in Spain in regions classified as "temperate, dry summer, warm summer" (Csb) under the Köppen-Geiger climate classification. (*) shows significant effects.

| MAINLAND (q=2) | | | | | | | |
|---|---|---|---|---|---|---|---|
| | | **Effect** | **Bias** | **Std. Err.** | **Lower CI** | **Upper CI** | |
| DIRECT | environmental conditions | -0.149 | 0.000 | 0.024 | -0.196 | -0.100 | * |
| | taxonomic div | -0.015 | 0.000 | 0.024 | -0.063 | 0.032 | |
| | phylogenetic div | 0.077 | 0.000 | 0.030 | 0.016 | 0.134 | * |
| | functional div | -0.004 | 0.000 | 0.026 | -0.054 | 0.048 | |
| | number of individuals | 0.222 | 0.000 | 0.025 | 0.171 | 0.271 | * |
| INDIRECT | environmental conditions | 0.029 | 0.000 | 0.008 | 0.016 | 0.045 | * |
| | non-native species | -0.006 | 0.000 | 0.003 | -0.012 | 0.000 | |
| | number of individuals | 0.002 | 0.000 | 0.003 | -0.002 | 0.008 | |
| TOTAL | environmental conditions | -0.120 | 0.000 | 0.022 | -0.163 | -0.076 | * |
| | non-native species | -0.006 | 0.000 | 0.003 | -0.012 | 0.000 | |
| | taxonomic div | -0.015 | 0.000 | 0.024 | -0.063 | 0.032 | |
| | phylogenetic div | 0.077 | 0.000 | 0.030 | 0.016 | 0.134 | * |
| | functional div | -0.004 | 0.000 | 0.026 | -0.054 | 0.048 | |
| | number of individuals | 0.224 | 0.000 | 0.025 | 0.174 | 0.272 | * |
| MEDIATORS | taxonomic div | -0.003 | 0.000 | 0.005 | -0.014 | 0.006 | |
| | phylogenetic div | 0.005 | 0.000 | 0.005 | -0.002 | 0.018 | |
| | functional div | -0.001 | 0.000 | 0.004 | -0.010 | 0.008 | |
| | number of individuals | 0.025 | 0.000 | 0.007 | 0.013 | 0.040 | * |

**Table S13.** Direct, indirect, total, and mediator effects of a structural equation model (SEM) exploring the influence of multifaceted diversity accounting for species abundances (q=2), environmental conditions, number of individuals (N), and the proportion of non-native species on productivity in **island** forests in Spain in regions classified as “temperate, dry summer, warm summer” (Csb) under the Köppen-Geiger climate classification. (*) shows significant effects.

| ISLAND (q=2) | | | | | | | |
|---|---|---|---|---|---|---|---|
| | | **Effect** | **Bias** | **Std. Err.** | **Lower CI** | **Upper CI** | |
| DIRECT | environmental conditions | -0.202 | -0.001 | 0.032 | -0.262 | -0.137 | * |
| | non-native species | 0.089 | -0.001 | 0.071 | -0.053 | 0.223 | |
| | phylogenetic div | 0.081 | -0.001 | 0.048 | -0.011 | 0.175 | |
| | number of individuals | 0.158 | 0.000 | 0.033 | 0.094 | 0.222 | * |
| INDIRECT | environmental conditions | -0.047 | 0.001 | 0.010 | -0.070 | -0.029 | * |
| | non-native species | 0.008 | 0.000 | 0.008 | 0.000 | 0.035 | |
| | number of individuals | 0.036 | -0.001 | 0.021 | -0.005 | 0.079 | |
| TOTAL | environmental conditions | -0.248 | 0.000 | 0.031 | -0.308 | -0.184 | * |
| | non-native species | 0.097 | 0.000 | 0.070 | -0.045 | 0.023 | |
| | phylogenetic div | 0.081 | -0.001 | 0.048 | -0.011 | 0.175 | |
| | number of individuals | 0.194 | -0.001 | 0.029 | 0.137 | 0.253 | * |
| MEDIATORS | phylogenetic div | 0.029 | 0.000 | 0.019 | -0.003 | 0.072 | |
| | number of individuals | -0.038 | 0.000 | 0.008 | -0.057 | -0.023 | * |
